# High-throughput identification of nuclear envelope protein interactions in *Schizosaccharomyces pombe* using an arrayed membrane yeast-two hybrid library

**DOI:** 10.1101/2020.07.29.227819

**Authors:** Joseph M. Varberg, Jennifer M. Gardner, Scott McCroskey, Snehabala Saravanan, William D. Bradford, Sue L. Jaspersen

## Abstract

The nuclear envelope (NE) contains a specialized set of integral membrane proteins that maintain nuclear shape and integrity and influence chromatin organization and gene expression. Advances in proteomics techniques and studies in model organisms have identified hundreds of proteins that localize to the NE. However, the function of many of these proteins at the NE remains unclear, in part due to a lack of understanding of the interactions that these proteins participate in at the NE membrane. To assist in the characterization of NE transmembrane protein interactions we developed an arrayed library of integral and peripheral membrane proteins in the fission yeast *Schizosaccharomyces pombe* for high-throughput screening using the split-ubiquitin based membrane yeast two hybrid system. We used this approach to characterize protein interactions for three conserved proteins that localize to the inner nuclear membrane: Cut11/Ndc1, Lem2, and Ima1/Samp1/NET5. Additionally, we determined how the interaction network for Cut11 is altered in canonical temperature-sensitive *cut11* mutants. This library and screening approach is readily applicable to characterizing the interactomes of integral membrane proteins localizing to various subcellular compartments.

## Introduction

The nuclear envelope (NE) is a double lipid bilayer that separates the nucleoplasm from the cytoplasm to allow for the compartmentalization of biological processes such as transcription and translation. Both the outer nuclear membrane (ONM) and inner nuclear membrane (INM) are enriched for specific nuclear envelope transmembrane proteins (NETs) that serve a wide variety of functions, including chromatin organization and regulation of gene expression, nuclear shape and dynamics, mechanosensation and signal transduction across the NE (1–3). NETs and their interacting partners at the nuclear periphery have been implicated in numerous human diseases collectively referred to as nuclear envelopathies and laminopathies (4–9). Studies of disease-associated NETs in humans have also identified striking patterns of cell and tissue specificity in NET expression and splicing as well as complex tissue-specific disease pathologies in NET mutants (10, 11). One proposed mechanism to explain these patterns of disease manifestation is that it is the result of disruption of interactions between NETs and their binding partners, which are also expressed in tissue-specific manner (10, 12). Accordingly, determining the mechanisms behind nuclear envelopathies requires an understanding of both the composition of the NE proteome, as well as the NET interactome.

Despite their clear clinical importance, the identification and functional characterization of NETs and their interacting proteins remains challenging. First, for decades, the list of NETs was small, restricted to a handful of abundant INM proteins. Advances in proteomics in the last two decades has expanded the list of candidate NETs to several hundred (13–17). However, determining which NETs are enriched at the INM and therefore have the potential to directly interact with the genome and participate in biological processes occurring in the nucleoplasm has traditionally required detailed studies of each NET using electron microscopy (EM). This low-throughput method has been essential to resolve the INM from the ONM, which are only separated by only 30-50 nm in most cells. Although super-resolution methods offer an alternative to EM, they too are low throughput. Smoyer et al. developed a high-throughput assay based on split-GFP to screen all known and predicted integral membrane proteins in *Saccharomyces cerevisiae* for access to the INM (18). The list of putative INM components included over 400 proteins, many of which also localize to other subcellular regions. Recent work also suggests that proteins are targeted to the INM for protein degradation through INM-specific quality control pathways (19). This adds yet another layer of complexity to the NET interactome. Additionally, the hydrophobic nature of many NETs makes their isolation and interactome characterization particularly difficult (20).

Significant insights into the function of many NETs has emerged from studies in model organisms including the budding yeast *S. cerevisiae* and the fission yeasts *Schizosaccharomyces pombe* and *Schizosaccharomyces japonicus*. For example, genetic and cell biological studies using these systems has identified conserved mechanisms for NETs in NE repair and quality control (21–24), regulation of nuclear size and shape (25–28) and active lipid metabolism at the INM (29). The founding member of the Sad1-Unc84 (SUN) domain-containing proteins, which play roles in chromosome organization, centrosome function and nuclear migration and positioning in many eukaryotes, was originally identified in a screen for yeast mutants defective in spindle formation (30). Sad1 is a component of the *S. pombe* spindle pole body (SPB), the yeast centrosome-equivalent organelle. Sad1 assists in the insertion of SPBs into the NE to facilitate the nucleation of the mitotic spindle microtubules (31, 32). In addition, Sad1 also facilitates centromere and telomere tethering in both mitosis and meiosis (33–36).

Like Sad1, Cut11 is also a component of the SPB involved in SPB insertion (32, 37); however, as the fission yeast ortholog of the conserved integral membrane protein Ndc1 it is perhaps best known for its conserved role in the NE tethering of nuclear pore complexes (NPCs) (38, 39). Ima1 is the *S. pombe* ortholog of the mammalian INM protein Samp1/NET5 that has conserved functions in nuclear organization (40, 41). It transiently localizes to the SPB early in mitosis in *S. pombe*, similar to its localization to the spindle in mammals (42). Lem2 is one of the two fission yeast Lap2-Emerin-Man1(LEM) domain-containing proteins (the second being Man1). Unlike Ima1 and Cut11, which transiently localize to the SPB during mitosis, Lem2 is found at the SPB throughout interphase and it is present at the INM during the entire yeast cell cycle (37, 41). Lem2 and Man1 play at least two roles at the INM: heterochromatin tethering (40, 43–46), and regulation of NE composition, integrity and structure (21, 41, 47–50). How these diverse functions are controlled is poorly understood in part because we lack sufficient knowledge of the interactome for these NETs, even in model systems such as yeast.

The membrane yeast two-hybrid (MTYH) technology allows for the identification of interactions between full-length integral membrane proteins heterologously expressed in *S. cerevisiae* (51). Previously, we adapted this technology to study the interactome of Ndc1 in *S. cerevisiae* (52). Not only did we identify novel Ndc1 interacting proteins, but the MYTH approach enabled us to test *ndc1* mutant alleles for defects in binding to various substrates and better understand the phenotypic differences we observed for these alleles. Here, we have expanded this approach to study fission yeast NE membrane proteins using a newly developed library of 1037 *S. pombe* MYTH prey constructs. Using this library, we performed high-throughput screening to identify interactions for three highly conserved INM proteins of diverse structure and function: Cut11, Lem2 and Ima1. Additionally, we determined how canonical alleles of *cut11* alter the Cut11 interactome to better understand its role at the SPB and NPC in fission yeast. This library is a new resource for the *S. pombe* community to assist in the characterization of integral membrane protein interactions.

## Results

### Generation of an arrayed *S. pombe* MYTH prey library

MYTH was first described by Stagljar et al in 1998 (53) and has been used to identify interactions between integral membrane proteins from multiple species (54–58). In our system, membrane bait proteins of interest were fused to a C-terminal fragment of ubiquitin (Cub) and a transcription factor (TF) moiety containing both the *E. coli* LexA DNA-binding domain and the herpes simplex virus VP16 activation domain (Figure1A). Interaction of the bait-Cub fusion protein with prey proteins fused to the N-terminus of ubiquitin (Nub) reconstitutes a pseudoubiquitin molecule at the membrane, which is recognized and processed by deubiquitinating proteases, releasing the TF reporter to activate the expression of *HIS3* and *ADE2* reporter genes.

To facilitate high-throughput screening and analysis of integral membrane bait proteins in fission yeast, we generated an arrayed library of *S. pombe* prey proteins fused to Nub (Figure 1A-C) (Table S1). We initially targeted approximately 1300 proteins including all proteins with one or more known or predicted transmembrane domain. A total of 1037 unique prey constructs were successfully generated, including 81%(773/946) of all putative integral membrane proteins. The remaining 264 prey include soluble and peripheral membrane proteins with annotated functions at a variety of subcellular locations including the NE, the endoplasmic reticulum (ER) and the mitochondrion (Figure 1B). The arrayed prey library approach allows us to rapidly assess pairwise interactions with hundreds of individual prey proteins in a single assay. Since the position of each prey protein is known, quantification of colony growth and downstream data analysis is streamlined as compared to alternative approaches that require recovery of plasmid DNA and sequencing to identify positive interactions (51).

**Fig. 1.**
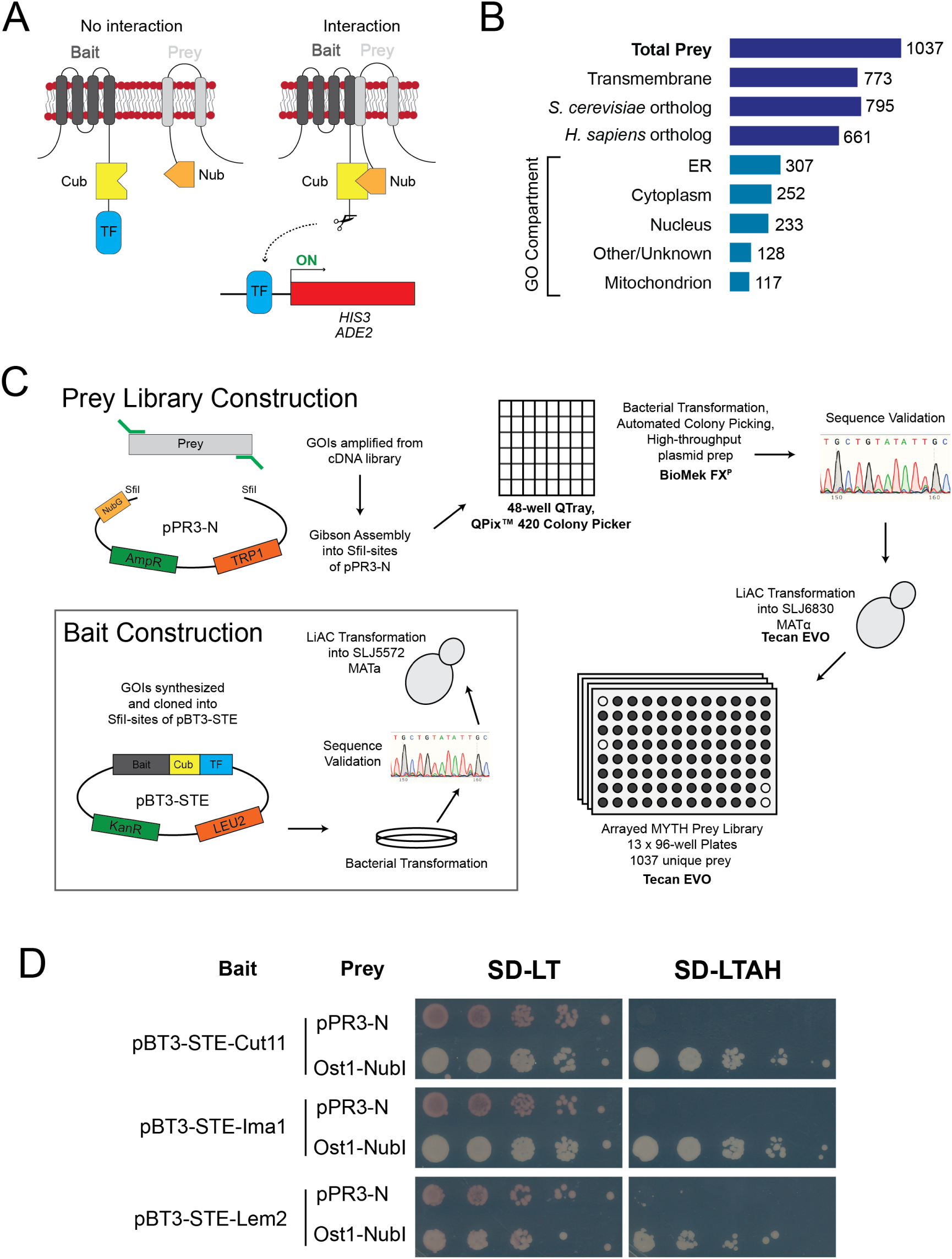
Generation of *S. pombe* MYTH prey library. A) Schematic of the split-ubiquitin MYTH approach. Full-length integral membrane bait proteins are fused to the C-terminus of ubiquitin (Cub) and a LexA-VP16 transcription factor reporter (TF). Upon interaction with a prey protein fused to N-terminus of ubiquitin (Nub), the ubiquitin molecule is reconstituted and cleaved by proteases to release TF for expression of *HIS3* and *ADE2* reporter genes. B) Diagram of *S. pombe* MYTH prey library composition by protein feature, conservation status and GO Compartment ID. C) Schematic of prey and bait construction approach. See Materials and Methods for detailed description. D) Validation of INM bait proteins for MYTH. Expression of both bait and prey plasmids is confirmed by growth on media lacking leucine (L) and tryptophan (T) (SD-LT). Cell growth on selective media further lacking both adenine (A) and histidine (H) (SD-LTAH) is only observed when each bait is co-expressed with a positive control Ost1-NubI prey protein.

### Bait selection, validation and library screening

To demonstrate the utility of the MYTH prey library we next sought to characterize the interactions for a collection of integral membrane proteins known to localize to the INM. Based on their conserved roles at the INM throughout eukaryotes, we focused on the SUN-domain protein Sad1, the nucleoporin (Nup) Cut11, the LEM-domain proteins Lem2 and Man1, and the Samp1/NET5 ortholog Ima1.

Strains expressing the INM baits were tested to confirm that the bait-Cub fusion proteins were expressed and that the growth of these strains on selective media was dependent upon prey interaction. Strains co-expressing individual baits with the empty prey plasmid (pPR3-N) fail to grow on selective media, while co-expression with a positive-control prey containing the Nub fragment that retains its affinity for Cub (Ost1-NubI) reconstitutes the pseudoubiquitin molecule and drives expression of the *HIS3* and *ADE2* reporter genes. Cut11, Ima1 and Lem2 C-terminal bait constructs (pBT3-STE) all passed these initial quality control assays (Figure 1D). Strains expressing Sad1 or Man1 baits were either not expressed or autoactivated (N-Sad1) (Figure S1). We therefore proceeded with screening Cut11, Ima1 and Lem2 against the prey library. As each localize to distinct subregions of the INM and do not contain similar functional domains, comparison of hits across baits would enable us to identify specific and non-specific interactions. All three baits were screened against the MYTH prey library simultaneously to avoid variability in media composition and screening conditions, al-lowing for direct comparison of colony size across baits.

To screen the three baits of interest against the MYTH library, strains expressing each bait were crossed with the MYTH prey library in a high-throughput fashion (Figure 2A). Positive interactions between bait-prey pairs were determined in a semi-automated fashion using a manually defined colony density threshold (*see Materials and Methods*), with larger densities representing stronger interactions between bait and prey pairs (Figure S2A). This approach allowed for the examination of thousands of pair-wise interactions in approximately two weeks from initiation of cultures to completion of analysis.

**Fig. 2.**
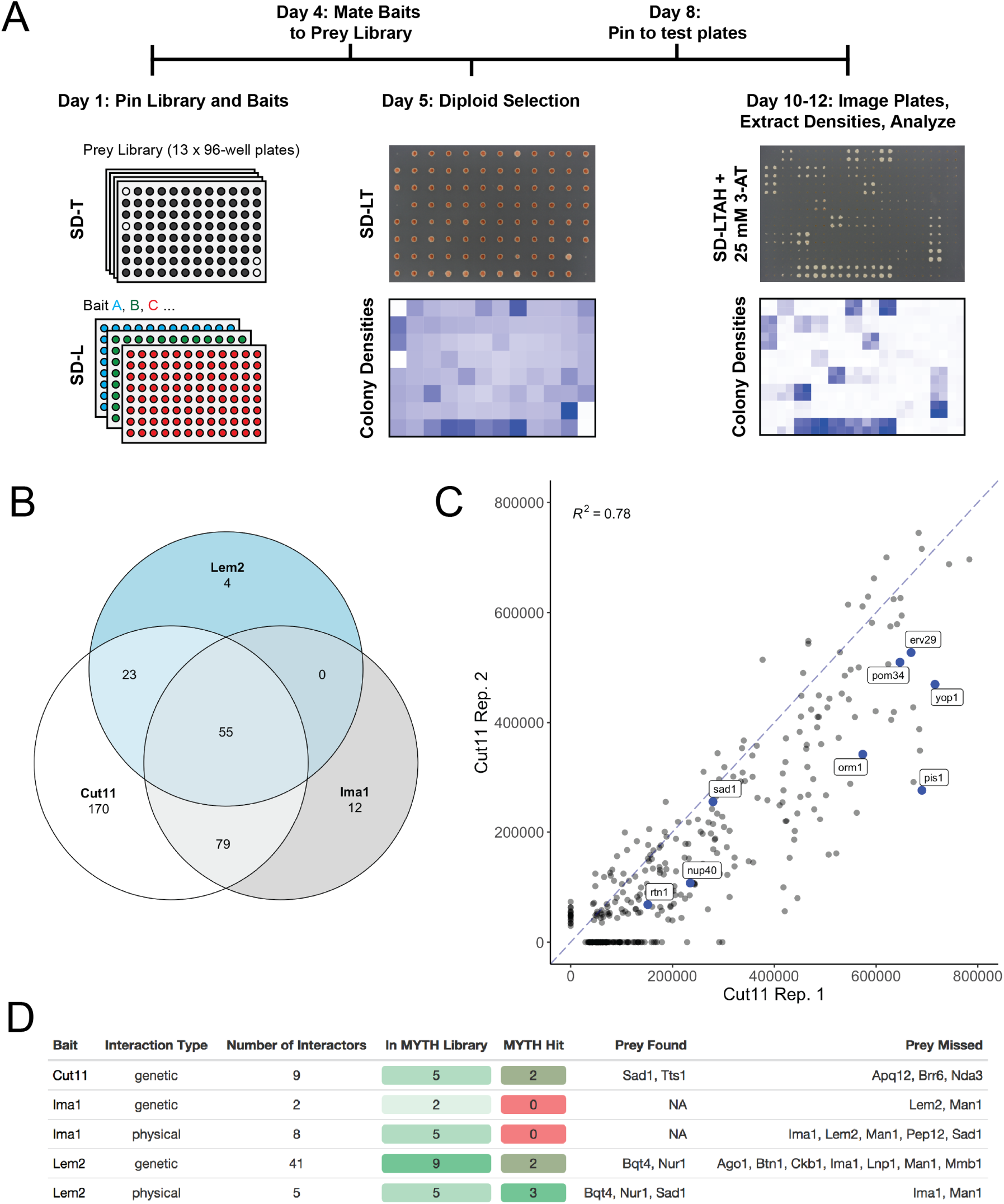
Screening for INM protein interactions using MYTH library. A) Overview of MYTH library screening. Strains expressing the MYTH bait and prey are grown on PlusPlates, mated and diploids are selected by growth on SD-LT media in 96-spot format. Positive bait-prey interactions are identified by monitoring colony growth on selective media (SD-LTAH) supplemented with 3-AT. Colony densities are extracted and prey identity is assigned based on known prey libray plate layout in an automated fashion. B) Venn diagram illustrating the number of shared or unique interacting prey proteins identified for each bait. C) Comparison of colony densities identified in independent Cut11 bait screen, showing high reproducibility in strength and identity of interactions identified by MYTH. Proteins known to interact with Ndc1 in *S. cerevisiae* are highlighted and labeled. D) Known genetic and physical interactors for each bait were examined to determine whether they were present in our prey library, and if they were identified as a positive hit in our screens.

Of the 1037 unique prey in the library, a total of 343 were identified as hits for at least one of the three INM baits that were screened (Figure 2B) (Table S2). We first examined whether our screen identified known physical interactions for each of our baits. Three of the five known Lem2 interacting proteins were identified as Lem2 hits in our screen (Bqt4, Sad1, Nur1, Figure 2D). Although no physical interactions have been reported for Cut11 in *S. pombe*, our MYTH screen identified 75% (9/12) of the prey that are orthologs of proteins reported to physically interact with Ndc1 in *S. cerevisiae* and were in our prey library (Figure 2C). In contrast, our screen failed to detect any of the five reported physical interactors for Ima1 that were present in our library. We noted that Man1 was not functional as a bait or prey, as the prey was not found to interact with Lem2 or Ima1, despite previously being identified as an interactor for both (41). We were unable to generate a functional Sad1 bait, however, the Sad1 prey construct was functional and recapitulated the known interaction with Lem2 (41, 59). Comparison of two independent screens using the same bait (Cut11) showed a high level of reproducibility in both the identity and strength of the prey interactions (Figure 2C). Together, these results demonstrate that our MYTH screen identified many known interactions with a high level of reproducibility; however, some proteins are incompatible with the MYTH system. This is likely due to defects in protein stability or folding, localization, or membrane topology introduced by expression as a bait or prey fusion protein.

### Comparison of interactions identified for each bait

We next performed pairwise comparisons between each bait to identify interactions that were common and those that were unique or enriched for specific baits. The strong positive correlation observed between replicates was not observed in comparisons between baits, and only 55 prey (16% of all hits) displayed interactions with all three baits (Figure 2B; Figure 3). A subset of these common interactors were among the strongest interactions identified for all three baits and likely represent non-specific interactions resulting in false positives. However, this group also included prey that displayed significant preferences for certain baits. For example, the trans-membrane Nup Pom34, which forms a complex with Ndc1 in *S. cerevisiae* (60), interacted with all three baits yet showed a 5-7-times stronger interaction with Cut11 compared to Lem2 and Ima1 (Figure 3D). GO term enrichment analysis of the hits that were uniquely enriched (> 2-fold increased over other baits) for each bait failed to identify any functional enrichment for the interacting prey. We therefore focused our analysis on prey that shared localization or predicted function with each bait, as well as those interactions that were unique or significantly enriched. Interactions of interest for each bait are discussed below.

**Fig. 3.**
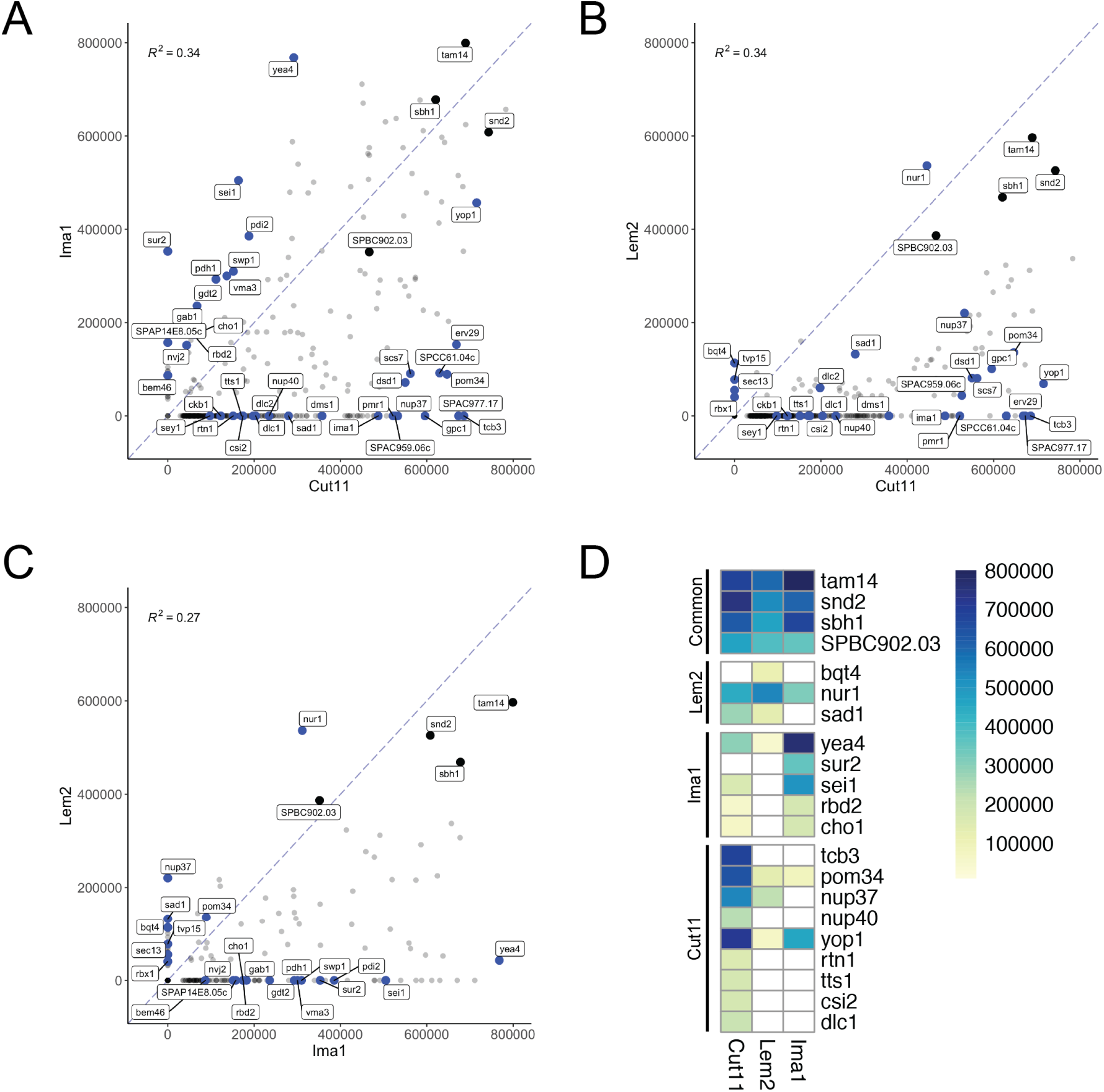
Comparison of INM bait interactomes. A-C) Scatter plots showing pairwise comparisons of average colony density values for prey identified as positive interactors for at least one bait protein in our screen. Prey that have a similar strength of interaction with both baits align on the diagonal dashed line. Prey of interest for each bait are annotated in blue, while four prey that were among the top interactions for all three baits are annotated in black. Pearson correlation coefficients are listed for each comparison. D) Heat map representation of select prey of interest that are discussed in text. Prey that showed no interaction with each bait are shown as white boxes.

### Lem2 MYTH Interactors

Lem2 is a conserved integral membrane protein that serves multiple functions at the INM. A total of 82 prey interacted with Lem2, of which only four were unique interactions (Bqt4, Sec13, Tvp15, Rbx1). Three of these unique interactions were very weak; however, we observed a strong unique interaction with the INM protein Bqt4, which anchors telomeres to the NE and binds directly to Lem2 to retain it at the NE (61, 62). Although not unique for Lem2, the Lem2-interacting protein Nur1 (43, 63, 64) was identified as the second strongest Lem2 interactor and was enriched 2-3-fold relative to Cut11 and Ima1. Identification of Bqt4 and Nur1 is consistent with the role of Lem2, Bqt4 and Nur1 in silencing and retention of centromeres and telomeres at the nuclear periphery (43, 47). Lem2 interacts directly with Bqt4 and Nur1 through its N-terminal LEM domain (44), providing evidence that MYTH is able to identify bona fide binding proteins for NETs.

We further examined Lem2 hits in an attempt to gain insight into proteins that may assist in its functions in heterochromatic gene silencing, telomere positioning and centromere tethering at the SPB. Unfortunately, many of the heterochromatin-associated factors that showed genetic interactions with Lem2 (44) were absent from our prey library, and none of the 14 heterochromatin factors present in our library interacted with Lem2 (Ago1, Stc1, Raf2, Pcu4, Chp1, Swi6, Clr4, Lem2, Man1, Nup85, Mmi1, Red1, Shf1, Brl1). Of the factors associated with telomere localization present in our library, Sad1, Bqt3 and Bqt4 were found to interact with Lem2 (Figure S2C). The only strong interaction we observed with known SPB-localized proteins was with Sad1, which has previously been shown to interact with Lem2 (41, 59). Thus, despite confirming known interactions with Sad1, Bqt4 and Nur1, interactions that physically connect Lem2 with components of the heterochromatin machinery still remain elusive (65).

Lastly, we examined the Lem2 interacting prey for factors potentially associated with NPC quality control at the NE. In *S. pombe*, Lem2 is required for maintaining NE morphology and regulating membrane flow from the ER into the NE to control nuclear size (47–49). However, a direct role for Lem2 in NPC quality control has not been shown in *S. pombe*, and Lem2 is not known to interact directly with any components of the NPC or the ESCRT (endosomal sorting complexes required for transport) machinery that facilitate NE/NPC quality control in other organisms. In budding yeast, the Lem2 ortholog Heh1/Src1 binds with luminal domains of the trans-membrane Nup Pom152 and displays genetic interactions with Nups in multiple subcomplexes (66). This interaction with the NPC is thought to allow Heh1/ Lem2 to recruit the ESCRT component Chm7 to regions of the NE that contain defects in NPC assembly to ensure NE compartmentalization is maintained (22, 23). Although we were unable to generate a functional *S. pombe* Pom152 MTYH prey, Lem2 interacted with the transmembrane Nup Pom34 (Figure 4D). We also observed an interaction between Lem2 and the inner ring component Nup37, which was recently found to associate with Lem2 in mammalian cells (67). Our results, in combination with the conserved interaction between Lem2 and Chm7 in *S. pombe* (21), their physical interaction in *S. japonicus* (46) and genetic interactions between Nups and the ESCRT machinery (68) suggests that a similar mechanism could be conserved.

**Fig. 4.**
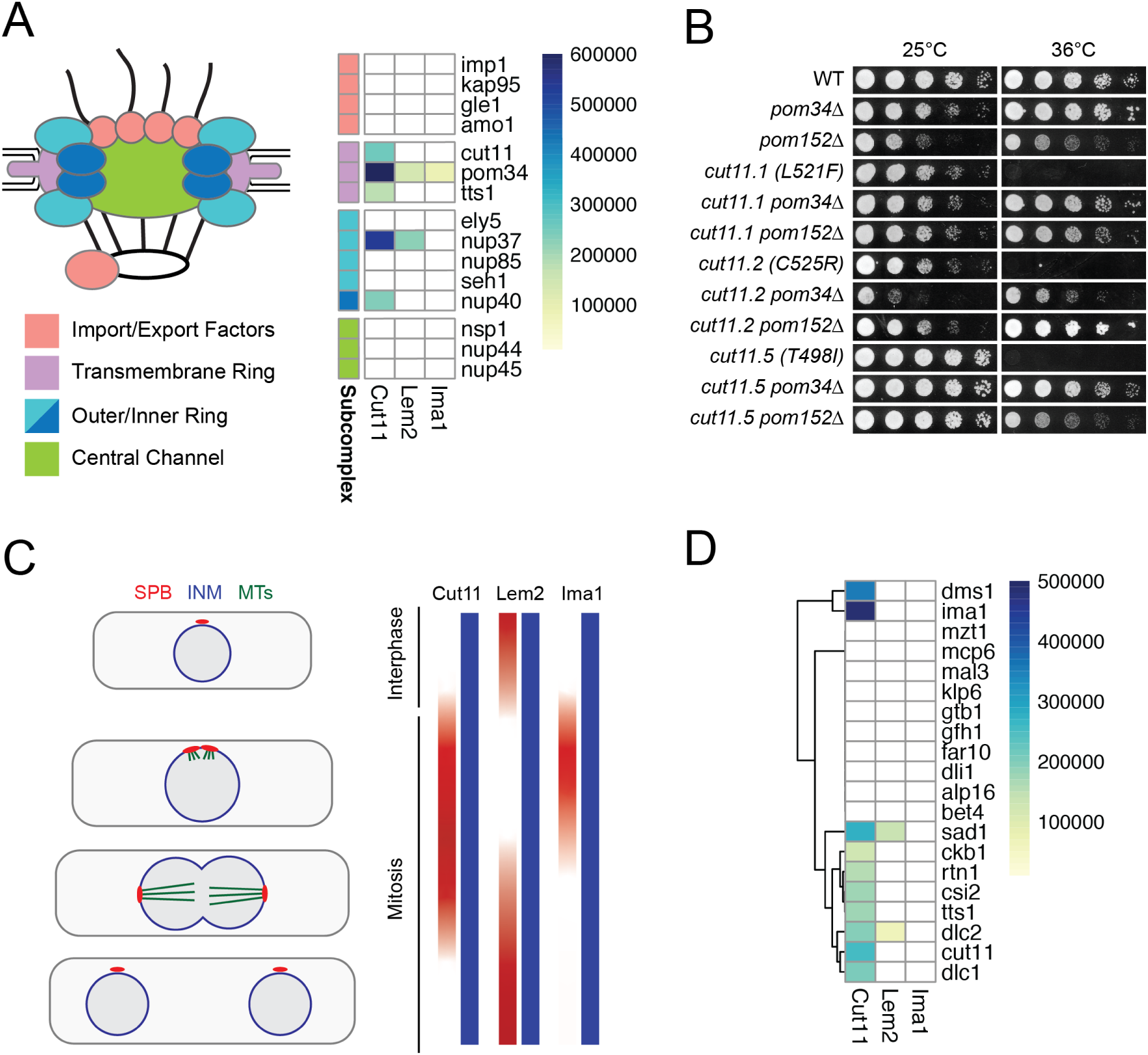
Interactions identified with NPC and SPB components. A) Cartoon representation of NPC structure (*left*) with subcomplexes colored to match groupings shown in heat map (*right*). B) Dilution assays assessing growth of *cut11* wild-type or temperature-sensitive alleles at permissive (25°C) and restrictive (36°C) temperatures. Deletion of either Pom34 or Pom152 rescues the temperature sensitivity of all *cut11-ts* alleles. C) Cut11, Lem2 and Ima1 each show enrichment at the SPB but at different stages of the cell cycle. Lem2 localizes to the SPB throughout interphase but is absent during early stages of mitosis. In contrast, both Cut11 and Ima1 localize to the SPB during mitotic entry during the period where Lem2 is absent. All three proteins also localize throughout the INM during the entire cell cycle. D) Heat map of interactions with prey proteins that are components of the SPB (GO:0005816).

### Ima1 MYTH Interactors

Ima1 is the fission yeast ortholog of the mammalian INM protein Samp1/NET5 (14, 42), for which there is no apparent ortholog in *S. cerevisiae*. Samp1 influences the distribution of other INM proteins, including the Sad1 ortholog Sun1, is required for centrosome positioning and tethering to the NE during interphase, and localizes to transmembrane actin-associated nuclear (TAN) lines where it stabilizes SUN-domain containing linker of nucleoskeleton and cytoskeleton (LINC) complexes to promote nuclear movement (42, 69, 70). In *S. pombe*, Ima1 localizes to distinct sub-regions of the NE where it interacts with specific heterochromatic regions of the genome (40). Despite enrichment of Ima1 near the central core regions of the centromeres, Ima1 is not required for centromere tethering at the SPB (41, 71). The function of Ima1 in *S. pombe* is unclear, though it appears to share a redundant role in nuclear membrane morphology and structure with the LEM-domain proteins Lem2 and Man1 (41). Therefore, identification of Ima1 interactors, particularly novel binding partners, would shed light on its function and guide future functional studies.

Ima1 has eight reported physical interactions, five of which are present in the MYTH prey library (Ima1, Lem2, Man1, Pep12, Sad1). Unfortunately, none of these interactors were identified as Ima1 hits in our screen. While this could be due to expression or functionality of either the bait or prey constructs, the fact that we identified the Ima1 and Sad1 prey as hits in our Cut11 and/or Lem2 screen makes this unlikely. It is possible that Ima1 interactions with these proteins may be regulated in a manner not recapitulated in the heterologous system.

In both fission yeast and mammals, Ima1/Samp1 displays cell cycle dependent changes in its localization from the INM to the SPB/centrosome-associated membrane. Given this conserved, transient recruitment of Samp1/Ima1 to the centrosome during mitosis, we next examined the list of Ima1 interactors for factors known to localize to the SPB. Although we did not observe any interactions between the Ima1 bait and SPB-localized prey, we noted that Ima1 prey showed a specific and strong interaction with the Cut11 bait (Figure 4D). Ima1 localizes to the SPB specifically during the early stages of mitosis, temporally overlapping with the kinetics of Cut11 recruitment to the SPB during SPB insertion (Figure 4C) (37, 41). It is thus possible that Ima1 localization to the SPB is facilitated by its interaction with Cut11, although its interaction with Sad1 may also contribute.

We next examined the twelve interactors that were uniquely identified for Ima1. The strongest of these interactions was with Sur2, a conserved sphingolipid metabolic enzyme with bi-functionality as a sphingosine hydroxylase and Δ 4-desaturase (72–75). While a functional role for Sur2 at the INM remains to be explored in fission yeast, Sur2 was found to have access to the INM in *S. cerevisiae* (18), and the mammalian sphingolipid hydroxylase Smpd4 is enriched at the NE and physically interacts with components of the NPC (76, 77). Further, Sur2 substrates, including the long-chain base dihydrosphingosine, are enriched at the nuclear membrane where they play a key role in maintaining nuclear morphology in both yeast and mammalian cells (78). This suggests that an interaction with Ima1 may retain a pool of Sur2 at the INM in *S. pombe* where it could influence properties of the NE membrane by controlling local sphingolipid content. Ima1-enriched prey included other factors implicated in lipid biogenesis and membrane organization. For example, Ima1 had a strong interaction with the seipin ortholog Sei1, which has a conserved role in lipid droplet biogenesis at the ER and stabilizes lipid droplet-ER contact sites (79). Recently, Sei1 was found to localize to the INM in budding yeast where it is required for the formation of membrane bridges between the INM and nuclear lipid droplets (29). We also observed an interaction with Nvj2, a lipid-binding protein that localizes to membrane contact sites between the nucleus and vacuole and forms membrane bridges between the ER and Golgi network to alleviate high levels of ceramides through nonvesicular transport (80, 81). Although this function in ceramide transport was not attributable to the perinuclear ER pool of Nvj2, it is possible that similar mechanisms exist to transport ceramides out of the NE, and its interaction with Ima1 presents additional evidence linking Ima1 with factors that may influence membrane composition at the INM. Other Ima1 interactors implicated in lipid metabolism include the rhomboid protease Rbd2, which cleaves and activates the sterol regulatory element-binding protein (SREBP) transcription factor Sre1 and is present at the INM in *S. cerevisiae* (18, 82, 83) and the phosphatidylcholine synthesis protein Cho1 (84). The remainder of the Ima1-enriched interacting prey are largely uncharacterized proteins with a variety of predicted functions including ER-associated protein modifications and vesicular transport, three of which (Yea4, Bem46, SPAP14D8.05c) physically interact with the INM protein Man1 (59). As many of these factors are conserved in humans, further characterization of Ima1 and its interacting proteins in fission yeast will provide insight into their potential conserved functions at the nuclear envelope.

### Cut11 MYTH Interactors

Cut11/Ndc1 is a transmembrane Nup conserved between yeast and vertebrates and is required for NPC assembly (39, 85, 86). In budding yeast Ndc1 has genetic and physical interactions with the transmembrane Nups Pom34 and Pom152 (52, 60). These interactions are required for NPC assembly and influence the distribution of Ndc1 in the NE by competing with the SUN-domain protein Mps3 for a shared binding site on Ndc1 (52). We previously used the MYTH system to show that deletion of Pom152 rescues *ndc1* temperature sensitive (ts) alleles by increasing Ndc1-Mps3 interactions to promote Ndc1 localization at the SPB (52). Although we were unable to generate a functional *S. pombe* Pom152 prey construct, we observed a strong interaction between Cut11 and Pom34 from our MYTH screen (Figure 4A). Further, deletion of either Pom34 or Pom152 rescued growth of each of the three causative mutations identified in *cut11-ts* strains (L521F (*cut11*.*1*), C525R (*cut11*.*2/3/4*) and T498I (*cut11*.*5/6*), Figure 4B) (87). The conservation of physical and genetic interactions between Ndc1/Cut11 and the Poms suggests that similar mechanisms controlling Cut11 localization and function in the NE may be conserved in fission yeast.

In addition to the Poms, Ndc1 also interacts with the structural nucleoporins Nup53 and Nup59 and the ER membrane-bending proteins Rtn1 and Yop1 to induce and stabilize membrane curvature that occurs during NPC assembly (60, 88–90). Similar interactions between Ndc1 and Nup53 occur in *Xenopus* (91) and vertebrate systems (86). Many of these factors were present in our prey library and were identified as strong, often unique, interactors for Cut11, including the *S. pombe* Nup53/59 ortholog Nup40, Rtn1, and Yop1 (Figure 3,4A). Additionally, both Yop1 interacting proteins, Yip1 (SPCC61.04c) and Sey1, which work together to form highly curved ER membrane tubules (92–94), facilitate lipid transfer between membranes and organelles (95), and influence NPC organization (90) interacted with Cut11 (Figure 3). Cut11 interacted with a specific subset of NPC components, as many Nups in our prey library showed no interaction with Cut11, including the structural Nups Ely5 and Nup85, and the FG-Nups Nsp1, Nup44 and Nup45 (Figure 4A). We did observe an interaction between Cut11 and Nup37, a structural Nup that is conserved in vertebrates but missing in *S. cerevisiae*. This suggests that a direct interaction between Cut11 and Nup37 may help to anchor the Nup107-160 subcomplex to the pore membrane in *S. pombe*. This interaction may also promote recruitment of the Nup107-160 complex to sites of NPC assembly, similar to the POM121-mediated recruitment reported in metazoans (96). Our screen also revealed an interaction between Cut11 and Tts1, a conserved reticulon-binding protein that has functions in membrane-shaping at the ER and NE (87, 97, 98). This was particularly exciting, as Tts1 displays genetic interactions with Cut11 in *S. pombe*, is implicated in NE remodeling during SPB insertion and in controlling NPC distribution during mitotic NE expansion (87) and co-purifies with fission yeast NPCs (64).

In addition to its function at the NPC, Cut11 is also required for SPB insertion and tethering within the NE during mitosis (32, 37). In budding yeast, the SUN-domain protein Mps3 controls the distribution of the Cut11 ortholog Ndc1 in the NE and facilitates its recruitment to the SPB, where it forms a membrane ring structure alongside other members of the SPB Insertion Network (SPIN) (99–101). We have recently shown that Cut11 forms similar ring-like structures during SPB duplication and insertion in *S. pombe* (unpublished data); however, the mechanisms that regulate Cut11 distribution and localization at the SPB remain unknown. To identify potential Cut11 interactors at the SPB, we examined interactions with known SPB-associated proteins (Figure 4D). We observed a strong interaction with the Mps3 ortholog Sad1, providing the first evidence for a direct interaction between these two proteins in *S. pombe* and suggesting that Sad1 may similarly promote Cut11’s localization at the SPB.

Cut11 also interacted with Csi2, which localizes to the SPB and is implicated in bipolar spindle formation, microtubule dynamics and chromosome segregation (102). Also identified was Ckb1, the beta-subunit of casein kinase CK2, which phosphorylates the HP1 protein Swi6 to promote heterochromatic silencing at centromeres (103, 104). Interestingly, we also identified SPB components with meiosis-specific functions. For example, Cut11 interacted strongly with Dms1, which localizes to the SPB specifically during meiosis II and recruits Spo15 for meiotic SPB remodeling to ensure proper SPB number and function (105–107). Cut11 also interacted with the dynein light-chain proteins Dlc1 and Dlc2. Dlc1 localizes to the SPB throughout mitosis and meiosis and was shown to interact with the meiosis-specific SPB component Kms1 by traditional yeast two-hybrid screen (108). Dynein motors are required for the “horse-tail” movements of telomere-associated SPBs during meiosis (109–112) and promote chromosome segregation during mitosis (113) and meiosis (114, 115). Together, these findings identify multiple putative new interacting proteins for Cut11, including components of the NPC and SPB, and demonstrate that the use of a heterologous system for the MYTH screening allows for the identification of interactions that may occur throughout the life cycle.

### Application of MYTH to determine how mutations alter interaction networks

We previously used the MYTH system to identify proteins that interact with Ndc1 at the SPB and NPC and to characterize how these interactions are modulated in *ndc1* mutants (52). As we observed conservation of physical and genetic interactions with Cut11 and the orthologous NPC and SPB components in fission yeast (Figure 4), we next examined the effects of canonical *cut11-ts* alleles on Cut11 localization and interactions. The original *cut11*.*1* mutant allele, leucine 521 to phenylalanine (L521F), is defective in SPB insertion and bipolar spindle formation, but has not been associated with defects in NPC assembly (37). Interestingly, this allele was not suppressed by increased levels of Tts1, while all other known alleles were, including *cut11*.*2/3/4*, which all contain a mutation of cysteine 525 to arginine (C525R), and *cut11*.*5/6* that contain a mutation of threonine 498 to isoleucine (T498I). All three residues map to the C-terminal tail of Cut11 that faces the nucleoplasm/cytoplasm, with L521F being highly conserved (Figure 5A-B). Visualization of each of the mutant proteins C-terminally tagged with GFP showed that all were expressed at similar levels and localized to punctate structures throughout the NE in interphase and to two bright foci in dividing cells (Figure 5C). This suggests that although these mutants show decreased protein levels relative to wild-type even at the permissive temperature (25°C), they properly localize to both NPCs and SPBs similarly to wild-type Cut11.

**Fig. 5.**
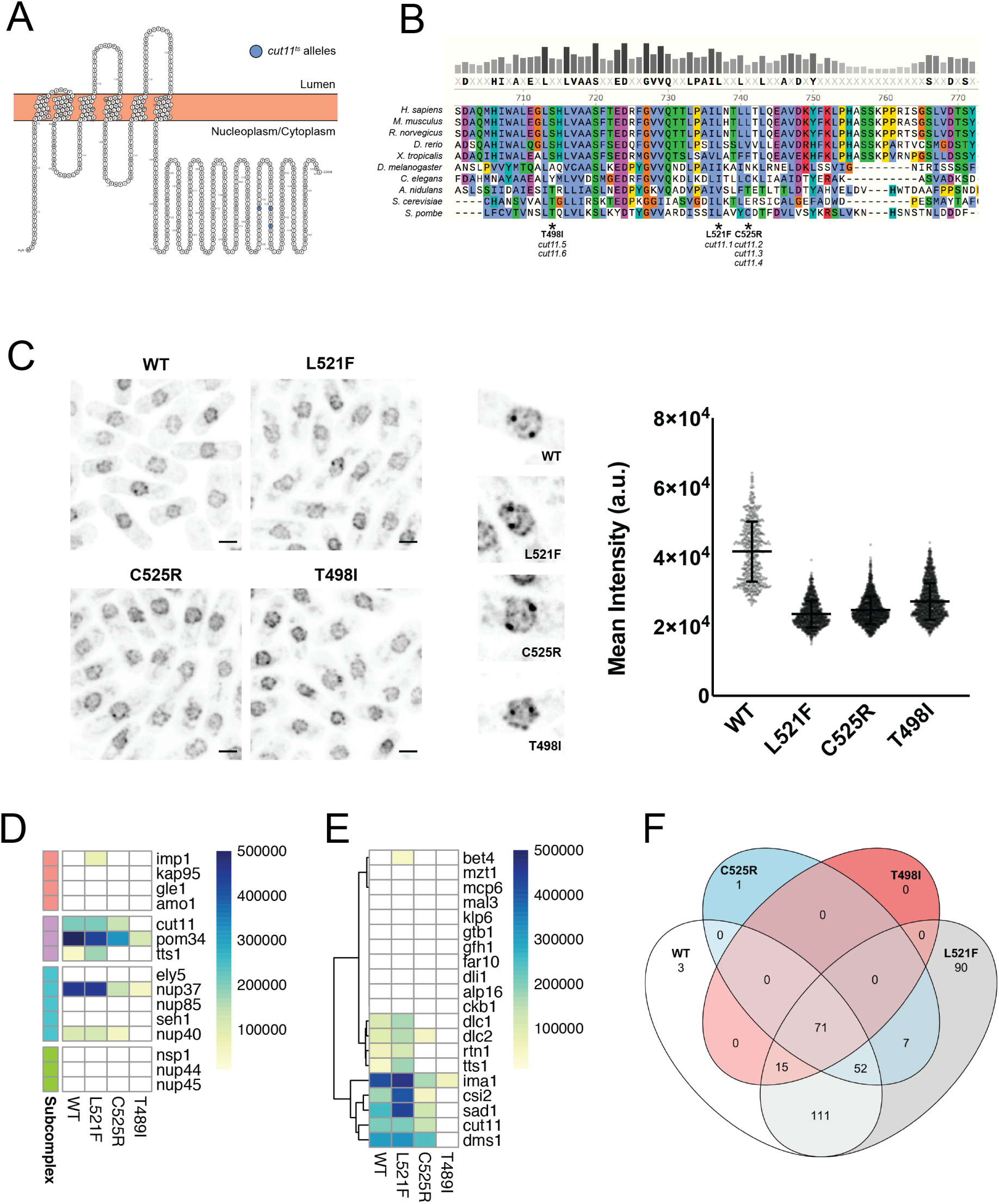
Effect of Cut11 mutations on the Cut11 interactome. A) Schematic of Cut11 topology in the NE, with mutations identified in *cut11-ts* alleles highlighted. All three mutations cluster in a similar region of the C-terminal portion of Cut11 that faces the nucleoplasm/cytoplasm (Image generated using Protter (http://wlab.ethz.ch/protter/start/)). B) C-terminal regions of Ndc1/Cut11 from multiple species aligned using ClustalOmega in SnapGene (v 5.1.4.1), with coloring based on amino acid properties and conservation. Causative mutations of each *cut11-ts* allele is annotated below alignment. C) (*left*) Representative images of Cut11 wild-type or mutant GFP fusion proteins, scale bar is 3 microns. (*center*) Images of individual mitotic nuclei showing recruitment of Cut11-GFP to the two SPBs (images show a 7.5×7.5 micron field of view). (*right*) Quantification of mean Cut11-GFP intensity values for individual nuclei in wild-type or mutant Cut11 strains. D-E) Heat maps of interactions between Cut11 baits and NPC (D) or SPB (E) components. The subcomplex color scheme in (D) is the same as used in Figure 4A. F) Venn diagram illustrating the overlap in interacting preys for each Cut11 bait protein.

To begin to understand the phenotypic differences reported for the *cut11-ts* mutants, we introduced the corresponding L521F, C525R or T498I mutations into the Cut11 MYTH bait. Expression of the mutant baits was confirmed (Figure S2D), and all three were screened against the MYTH prey library simultaneously with wild-type Cut11 to examine how these mutations altered protein interactions. Interestingly, the interactome of *cut11*.*1* (L521F) was similar to that of wild-type Cut11, with strong interactions with NPC components (Nup37, Pom34, Nup40, Cut11) and SPB components (Sad1, Ima1, Csi2) (Figure 5D-F). In contrast, both C525R and T498I mutants showed global alterations to their interactomes including reduced affinity for both NPC and SPB components. Inspection of the interaction with Tts1 showed an increase in binding for L521F compared to wild-type and a complete loss of binding for both C525R and T498I. Thus, genetic suppression by Tts1 overexpression can be linked to differences in the ability of different mutants to interact with Tts1. These results demonstrate that MYTH is a simple alternative tool to study the effect of mutations on protein interactions to provide additional insight into phenotypes observed in *S. pombe*.

## Discussion

Proteins that localize to the NE play critical roles in nuclear and chromatin organization, regulation of gene expression, lipid biosynthesis and membrane organization. To assist in the functional characterization of integral membrane proteins that localize to the NE, we developed an arrayed membrane yeast two-hybrid prey library. Screening of three known INM proteins against this library confirmed many previously reported interactions and identified novel interactions that are intriguing candidates for further mechanistic studies. This library and screening approach is immediately applicable to survey interactions that occur at other membranes and organelles in either a high-throughput or targeted approach. The MYTH system is also a valuable tool for determining how mutations of integral membrane proteins affect their protein interactions. By combining MYTH with genetic approaches available in *S. cerevisiae* and *S. pombe*, we have identified conserved interactions between Ndc1/Cut11 and components of the NPC and SPB that control its distribution in the NE. Characterization of the interaction profiles for canonical *cut11* alleles identified changes in interactions with putative Cut11 binding partners at the SPB and NPC. These studies also revealed a direct physical interaction between Cut11 and the conserved membrane protein Tts1. The observation that *cut11* alleles differentially alter interactions with Tts1 provides valuable new insight into the mechanisms by which Tts1 and Cut11 work together at the NE. In addition to its role in controlling NPC distribution during NE expansion, Tts1 deletion also exacerbated the spindle defects observed in *cut11-ts* mutants. The mechanism by which Tts1 exerts its function at the SPB remained unclear, as Tts1 does not localize to the SPB and is not required for localization of Cut11 to the SPB (87). It was therefore proposed that Tts1 likely promotes NE remodeling during SPB insertion by regulating membrane lipid composition or dynamics. Our data showing that mutants that do not bind to Tts1 (C525R and T498I) still localize to the SPB (Figure 5C) provides additional evidence that binding to Tts1 is not required for recruitment of Cut11 to the SPB or for SPB insertion.

Our results also allow us to address the curious observation that Tts1 overexpression rescues all *cut11-ts* alleles except for *cut11*.*1* (87). Our MYTH data show that the *cut11*.*1* L521F mutation increases binding between Cut11 and Tts1, while both C525R and T498I mutations prevent binding to Tts1. As Tts1 localizes to the INM and NPC but not the SPB (87), increased levels of Tts1 could act as a sink to retain Cut11 at these locations at the NE, potentially reducing the amount of Cut11 able to localize to the SPB to facilitate insertion. This impact of Tts1 overexpression is dependent upon its ability to bind to Cut11, and therefore is not observed in C525R or T498I mutants. This model also suggests that while Tts1 overexpression may promote changes to the membrane that are favorable for SPB insertion, this process requires Cut11 localization and function at the SPB.

A mechanism by which Tts1 overexpression competes with SPB components for Cut11 binding is similar to our data from budding yeast and fission yeast showing that *ndc1/cut11-ts* alleles can be rescued by reducing affinity for the NPC by disrupting interactions with the Pom nucleoporins (52) (Figure 4B). In budding yeast, distribution of Ndc1 between the NPC and SPB is facilitated by the SUN protein Mps3. Although we observe a conserved interaction with the *S. pombe* SUN protein Sad1, it is unclear whether Sad1 serves a similar function given that its localization at the INM is restrained to the regions near the SPB. It is likely that Sad1 serves a more passive role in *S. pombe*, acting as an anchor to retain Cut11 at the SPB but not actively shuttling Cut11 in the NE. In this scenario, recruitment of Cut11 to the SPB during mitosis may be controlled by altering the affinity for Cut11’s binding partners at the NPC or NE. Our data shows that changes within the C-terminus of Cut11 influence its protein interactions. Multiple residues in this region are phosphorylated in a cell-cycle dependent manner (116, 117) and could drive similar alterations to the Cut11 interactome to promote its recruitment to the SPB.

It also remains unclear how the C525R and T498I mutant proteins localize to the SPB, as many of the SPB interactions including Sad1 are significantly reduced or completely lost in these mutants. Both C525R and T498I are able to bind to Ima1, which localizes to the mitotic SPB with similar kinetics as Cut11 (Figure 4C). However, a direct role for Ima1 in Cut11 recruitment has not yet been reported. It is also likely that other SPB components that are not in our MYTH prey library, such as the KASH-protein Kms2 and the mitotic regulator Cut12, remain capable of binding and recruiting C525R and T498I mutant Cut11 proteins. The interactome data for the *cut11* alleles presented in this study will help guide future mechanistic studies exploring Cut11 function. MYTH may also be a helpful tool to identify mutations that will allow for separation of SPB and NPC function, which would be useful for studies characterizing the role of this conserved nucleoporin in NPC assembly and insertion into the NE.

## Methods

### A. Media preparation and yeast culture

Standard meth-ods were used for both *S. cerevisiae* (118) and *S. pombe* (119) transformation and colony selection. Synthetic drop-out (SD) media lacking the indicated amino acids was prepared by mixing 6.7 g yeast nitrogen base without amino acids with ammonium sulfate, 20 g dextrose (Sigma), 20 g Bacto Agar (VWR) and 0.5-1 g amino acid drop-out powder (Sunrise Scientific) in 1 L of water. Yeast extract with supplements (YES) media was prepared by mixing 5 g yeast extract, 30 g dextrose, 0.2 g each adenine, uracil, histidine, leucine and lysine, in 1 L of water. MYTH screens were performed in 96-or 384-well format on PlusPlates (Singer Instruments), at the temperatures indicated in the section below. Dilution assays to assess growth for MYTH bait quality control experiments were done using a serial 10-fold dilution series spotted onto control (SD-leu-trp) or selection media (SD-leu-trpade-his). Growth assays for *cut11-ts* rescue by Pom deletion were done using a 5-fold serial dilution series spotted onto YES agar plates incubated at permissive (25°C) or restrictive (36°C) temperatures.

### B. Generation of MYTH prey library

The coding sequence of each prey of interest was amplified from an *S. pombe* cDNA library (AS One International, Inc.) using KOD Hot Start DNA polymerase (Millipore Sigma), and reactions were cleaned up using the MagSi-DNA Clean paramagnetic beads (Amsbio). The amplicons were cloned into PCR-linearized pPR3-N prey plasmid (Dualsystems Biotech) at the SfiI sites using the NEBuilder HiFi DNA Assembly Kit (New England Biolabs), and were transformed into DH5*α* competent cells and plated onto 48-well Bioassay Qtrays (Molecular Devices). Automated colony picking was performed using a QPix 420 robotic colony picker (Molecular Devices), followed by high-throughput plasmid prep using a BioMek FXP liquid handling workstation (Beckman Coulter) and Sanger sequencing for insert validation. After sequencing validation, the prey plasmids were transformed into the MYTH prey reporter strain SLJ6830 (NMY61, Dualsystems Biotech) using traditional LiOAc protocol on a Freedom EVO automation platform (Tecan). Transformants were selected on SD-trp media, then picked and arrayed into a final format spanning thirteen 96-well plates using the Tecan EVO. In a similar fashion, MYTH bait coding sequences were gene synthesized (GenScript) and cloned into either the C-terminal (pBT3-STE) or N-terminal (pBT3-N) bait plasmids (pSJ1283 and pSJ1281). The bait plasmids were transformed into SLJ5572 (NMY51; Dualsystems Biotech) and transformants were selected on SD-leu plates.

### C. MYTH Screening

Liquid cultures for each bait were spotted onto SD-leu PlusPlates in 96-well format and 10 *µ*l volumes, and incubated for 2 days at 30°C. Similarly, the arrayed prey library was spotted in 96-well format on SD-trp PlusPlates. The bait and prey colonies were mated on YPD plates using a RoToR HDA-Robot (Singer Instruments Co. Ltd) and incubated overnight at 30°C. Diploids were selected by transferring the resulting colonies to SD-leu-trp media. Following diploid selection, each bait-prey combination was spotted in technical quadruplicate in 384-well format on SD-leu-trp-ade-his plates supplemented with 25 mM 3-aminotriazole (3-AT) to prevent leaky expression of the *HIS3* reporter gene (51, 120, 121).

### D. MYTH Analysis

Colony growth was monitored visually, and plates were imaged every 24 hours for 4 days. Colony densities were extracted using a custom FIJI/ImageJ plugin, and downstream analysis was conducted using RStudio (122). The code used for density extraction, data analysis and visualization are made available at http://www.stowers.org/research/publications/libpb-1540. Colony growth values obtained using this approach were nearly identical to measurements obtained using other tools designed for analysis of array-based high-throughput screens (123, 124) (Figure S2). To determine a density cutoff for assigning positive interactions, we manually assigned at least one hundred individual colonies to one of four categories based on strength of interaction: negative, weak, medium or strong. A density cutoff value equal to the 25th percentile of the “weak” interaction densities was chosen, and this threshold was applied to the entire screen. Interactions were considered “positive” if at least one-half of a prey’s spots on the test plate had a density value greater than the cutoff value. Values shown for positive interactors in Table S2 correspond to the averaged density values for each prey. GO term enrichment analysis was performed for prey that showed a 2-fold or greater enrichment for a specific bait, using LAGO (125) (https://go.princeton.edu/cgi-bin/LAGO), with the PomBase GOA annotations for biological processes, a p-value cutoff of 0.01 with Bonferroni correction, and the MYTH prey library as background.

### E. Microscopy and image analysis

Wild-type or *cut11-ts* strains were tagged at their endogenous locus with a C-terminal GFP tag. Cells were cultured in YES media at 25°C, and exponentially growing cells were collected and fixed for 20 minutes with 4% paraformaldehyde as previously described (32). Cells were resuspended in 1x PBS and imaged on a Nikon Ti2 wide-field microscope equipped with an Andor EM-CCD camera and an *α* Plan Apochromat 100x, 1.46 NA oil immersion objective. GFP fluorescence was excited with a 488nm (70mw) diode laser and collected with an ET535/30m emission filter. Data were acquired using Nikon Elements software (Nikon) with z-spacing of 300 nm covering a total volume of 6.3 *µ*m. All strains were imaged on the same day using constant exposure times, laser power and camera gain settings. Images were processed using FIJI/ImageJ (126) (National Institutes of Health, Bethesda, MD). To quantify Cut11-GFP levels at the NE, nuclear masks were created by automated thresholding of the Cut11-GFP signal. First, image stacks were maximum projected, background subtracted using a rolling ball radius of 20 pixels and blurred with a Gaussian blur filter. Thresholding was performed using the default algorithm in ImageJ, and nuclei were defined as particles with a size between 4 and 12 *µ*m^2^ and a circularity value of 0.3 or greater. The nuclear mask was then applied to a sum projection of the original image stack to extract mean intensity values for each nucleus. These values were plotted using the GraphPad Prism software package (v 8.4.3).

## ACKNOWLEDGEMENTS

We would like to thank Madeline Gogol and members of the Stowers Institute Molecular Biology Core Facility for assistance in generating the *S. pombe* prey library, and Jay Unruh and members of the Stowers Microscopy Core Facility for assistance in image analysis for colony density extraction. We also thank members of the Jaspersen lab for discussion and feedback on this manuscript. J.V. was funded by a Ruth L. Kirschstein NRSA Postdoctoral Fellowship (F32GM133096). The Jaspersen laboratory is supported by the Stowers Institute for Medical Research and by R01GM121443.

## DATA AVAILABILITY STATEMENT

Upon peer-reviewed publication, all strains generated for and used in this study will be made available upon reasonable request. Original data underlying this manuscript will be accessible at the Stowers Original Data Repository at http://www.stowers.org/research/publications/libpb-1540.

---

**Fig. S1.**
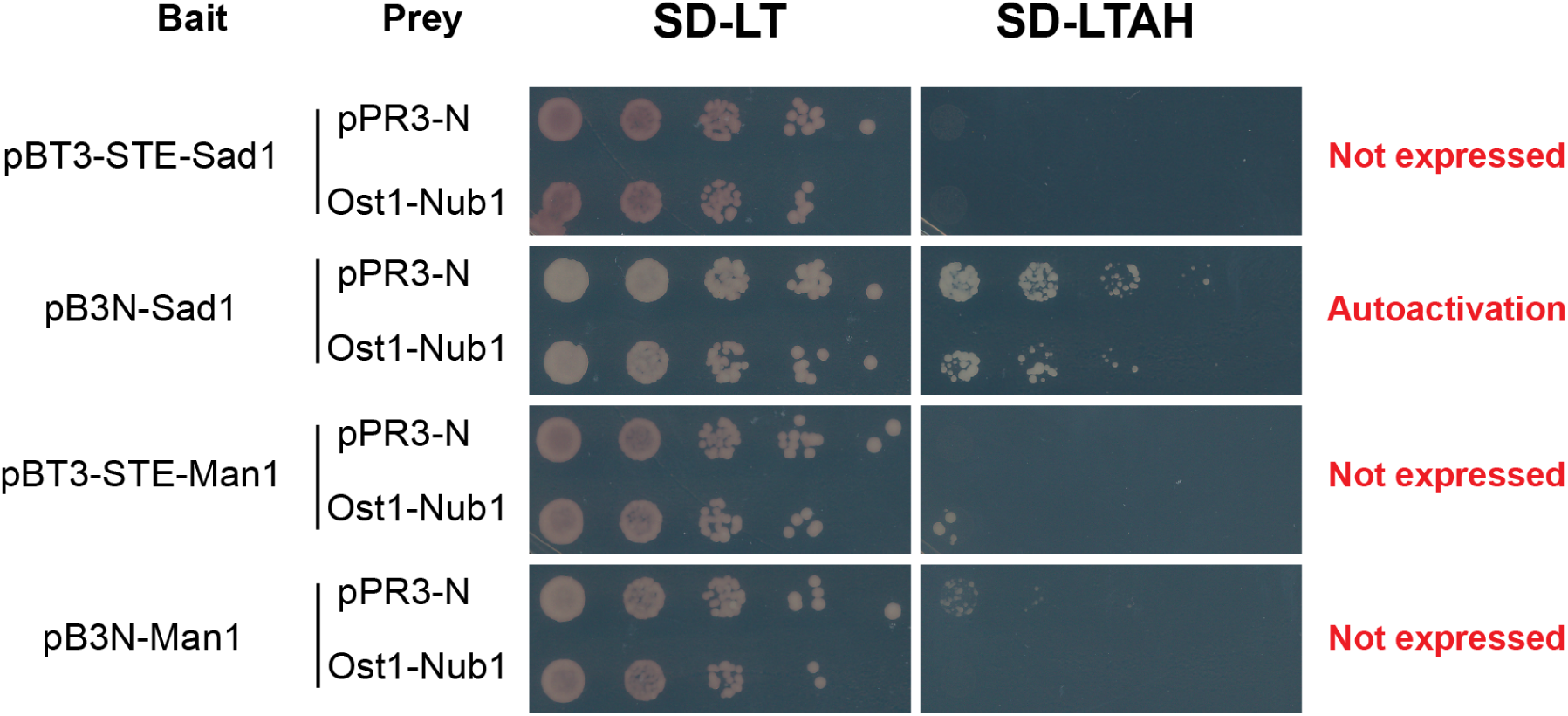
Quality control of MYTH baits and functional composition of prey library. Growth assays for MYTH baits that failed quality control screening. N-terminal Sad1 bait fusion protein showed growth on selection media when co-expressed with the empty MYTH prey plasmid (pPR3-N), indicating autoactivation. No growth was observed for C-terminal baits of Sad1 or Man, or for the N-terminal Man1 bait, when co-expressed with the positive control prey construct (Ost1-NubI), which contains a Nub fragment that retains its affinity for Cub.

**Fig. S2.**
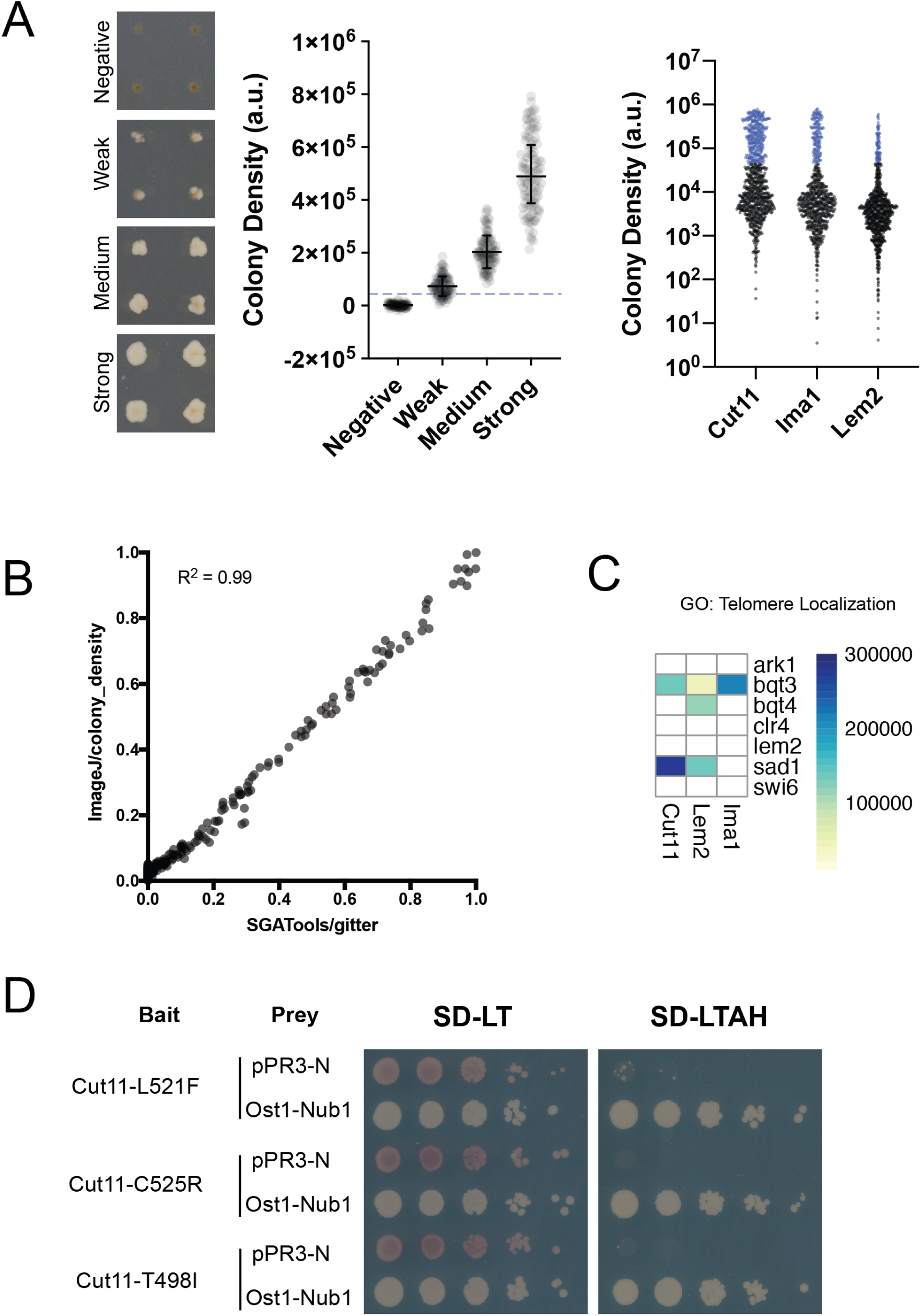
Criteria used to determine positive interactions A) (*left*) Representative images of MYTH colony growth after 4 days of growth on SD-leu-trp-ade-his + 25mM 3-AT test plates. At least 100 individual colonies were manually scored and binned into four categories: strong, medium, weak and negative. (*center*) A density cutoff was set at the 25th percentile value for “weak” colonies, depicted by dashed line. (*right*) Colony densities for each bait showing distribution of densities, with those falling above the threshold cutoff highlighted in blue. B) Comparison of normalized density values extracted using our custom ImageJ plugin with those available in the SGATools software package (http://sgatools.ccbr.utoronto.ca). C) Heat map of prey with GO annotation for telomere localization (GO:0034397). D) Growth assays confirming that Cut11 mutant baits are expressed and do not autoactivate.

## Supplemental Tables

**Table S1: *S. pombe* MYTH prey library** Each prey in libray is listed, with columns for Pombase StrainID, Gene, Product, presence of transmembrane (TM) domain, *S. cerevisiae* and *H. sapiens* orthologs, and the location of each protein as determined using a nucleoplasmic split-GFP reporter screen in *S. cerevisiae*. “INM” indicates observed localization to the inner nuclear membrane, “Nucleoplasmic” indicates soluble nuclear localization, “Negative” indicates no signal detected with nucleoplasmic reporter, and “Unknown” indicates that the protein was not included in the screen.

**Table S2: Interactors identified by MYTH screen** All 343 prey identified as a positive interactor for either Cut11, Ima1 or Lem2 bait proteins. Values represent averaged density values for all technical replicates for each prey.

**Table S3: Interactomes of wild-type and mutant Cut11** Prey identified as interactors for Cut11 bait proteins. Values represent averaged densities for all technical replicates for each prey. An additional column is provided showing fold-change relative to wild-type (WT) for each mutant bait (L521F, C525R and T498I).

## Notes

### Competing Interest Statement

The authors have declared no competing interest.

## Bibliography

1. Karim Mekhail and Danesh Moazed. The nuclear envelope in genome organization, expression and stability. Nature Reviews Molecular Cell Biology, 11(5):317–328, May 2010. ISSN 1471-0080. doi: 10.1038/nrm2894. Number: 5 Publisher: Nature Publishing Group.

2. Rosemarie Ungricht and Ulrike Kutay. Establishment of NE asymmetry—targeting of membrane proteins to the inner nuclear membrane. Current Opinion in Cell Biology, 34:135–141, June 2015. ISSN 0955-0674. doi: 10.1016/j.ceb.2015.04.005.

3. GW Gant Luxton and Daniel A Starr. KASHing up with the nucleus: novel functional roles of KASH proteins at the cytoplasmic surface of the nucleus. Current Opinion in Cell Biology, 28:69–75, June 2014. ISSN 0955-0674. doi: 10.1016/j.ceb.2014.03.002.

4. Raz Somech, Sigal Shaklai, Ninette Amariglio, Gideon Rechavi, and Amos J. Simon. Nuclear Envelopathies—Raising the Nuclear Veil. Pediatric Research, 57(7):8–15, May 2005. ISSN 1530-0447. doi: 10.1203/01.PDR.0000159566.54287.6C. Number: 7 Publisher: Nature Publishing Group.

5. Ya-Hui Chi, Zi-Jie Chen, and Kuan-Teh Jeang. The nuclear envelopathies and human diseases. Journal of Biomedical Science, 16(1):96, October 2009. ISSN 1423-0127. doi: 10.1186/1423-0127-16-96.

6. Alexandre Méjat and Tom Misteli. LINC complexes in health and disease. Nucleus, 1(1): 40–52, 2010. ISSN 1949-1034. doi: 10.4161/nucl.1.1.10530.

7. Alexandre Janin, Delphine Bauer, Francesca Ratti, Gilles Millat, and Alexandre Méjat. Nuclear envelopathies: a complex LINC between nuclear envelope and pathology. Orphanet Journal of Rare Diseases, 12(1):147, August 2017. ISSN 1750-1172. doi: 10.1186/s13023-017-0698-x.

8. Howard J. Worman, Cecilia Östlund, and Yuexia Wang. Diseases of the Nuclear Envelope. Cold Spring Harbor Perspectives in Biology, 2(2):a000760, February 2010. ISSN, 1943-0264. doi: 10.1101/cshperspect.a000760. Company: Cold Spring Harbor Laboratory Press Distributor: Cold Spring Harbor Laboratory Press Institution: Cold Spring Harbor Laboratory Press Label: Cold Spring Harbor Laboratory Press Publisher: Cold Spring Harbor Lab.

9. Patricia M. Davidson and Jan Lammerding. Broken nuclei – lamins, nuclear mechanics, and disease. Trends in Cell Biology, 24(4):247–256, April 2014. ISSN 0962-8924. doi: 10.1016/j.tcb.2013.11.004.

10. Nadia Korfali, Gavin S. Wilkie, Selene K. Swanson, Vlastimil Srsen, Jose de las Heras, Dzmitry G. Batrakou, Poonam Malik, Nikolaj Zuleger, Alastair R. W. Kerr, Laurence Florens, and Eric C. Schirmer. The nuclear envelope proteome differs notably between tissues. Nucleus, 3(6):552–564, November 2012. ISSN 1949-1034. doi: 10.4161/nucl.22257. Publisher: Taylor & Francis_eprint: https://doi.org/10.4161/nucl.22257.

11. Howard J Worman and Eric C Schirmer. Nuclear membrane diversity: underlying tissue-specific pathologies in disease? Current Opinion in Cell Biology, 34:101–112, June 2015. ISSN 0955-0674. doi: 10.1016/j.ceb.2015.06.003.

12. Jose I. de Las Heras, Peter Meinke, Dzmitry G. Batrakou, Vlastimil Srsen, Nikolaj Zuleger, Alastair Rw Kerr, and Eric C. Schirmer. Tissue specificity in the nuclear envelope supports its functional complexity. Nucleus (Austin, Tex.), 4(6):460–477, December 2013. ISSN 1949-1042. doi: 10.4161/nucl.26872.

13. Mathias Dreger, Luiza Bengtsson, Torsten Schöneberg, Henning Otto, and Ferdinand Hucho. Nuclear envelope proteomics: Novel integral membrane proteins of the inner nuclear membrane. Proceedings of the National Academy of Sciences, 98(21):11943–11948, October 2001. ISSN 0027-8424, 1091-6490. doi: 10.1073/pnas.211201898. Publisher: National Academy of Sciences Section: Biological Sciences.

14. Eric C. Schirmer, Laurence Florens, Tinglu Guan, John R. Yates, and Larry Gerace. Nuclear membrane proteins with potential disease links found by subtractive proteomics. Science (New York, N.Y.), 301(5638):1380–1382, September 2003. ISSN 1095-9203. doi: 10.1126/science.1088176.

15. Gavin S. Wilkie, Nadia Korfali, Selene K. Swanson, Poonam Malik, Vlastimil Srsen, Dzmitry G. Batrakou, Jose de las Heras, Nikolaj Zuleger, Alastair R. W. Kerr, Laurence Florens, and Eric C. Schirmer. Several Novel Nuclear Envelope Transmembrane Proteins Identified in Skeletal Muscle Have Cytoskeletal Associations. Molecular & Cellular Proteomics, 10(1), January 2011. ISSN 1535-9476, 1535-9484. doi: 10.1074/mcp.M110.003129. Publisher: American Society for Biochemistry and Molecular Biology Section: Research.

16. Nadia Korfali, Gavin S. Wilkie, Selene K. Swanson, Vlastimil Srsen, Dzmitry G. Batrakou, Elizabeth A. L. Fairley, Poonam Malik, Nikolaj Zuleger, Alexander Goncharevich, Jose de las Heras, David A. Kelly, Alastair R. W. Kerr, Laurence Florens, and Eric C. Schirmer. The Leukocyte Nuclear Envelope Proteome Varies with Cell Activation and Contains Novel Transmembrane Proteins That Affect Genome Architecture. Molecular & Cellular Proteomics, 9(12):2571–2585, December 2010. ISSN 1535-9476, 1535-9484. doi: 10.1074/mcp.M110.002915. Publisher: American Society for Biochemistry and Molecular Biology Section: Research.

17. Eric C. Schirmer and Larry Gerace. The nuclear membrane proteome: extending the envelope. Trends in Biochemical Sciences, 30(10):551–558, October 2005. ISSN 0968-0004. doi: 10.1016/j.tibs.2005.08.003.

18. Christine J. Smoyer, Santharam S. Katta, Jennifer M. Gardner, Lynn Stoltz, Scott Mc-Croskey, William D. Bradford, Melainia McClain, Sarah E. Smith, Brian D. Slaughter, Jay R. Unruh, and Sue L. Jaspersen. Analysis of membrane proteins localizing to the inner nuclear envelope in living cells. Journal of Cell Biology, 215(4):575–590, November 2016. ISSN 0021-9525. doi: 10.1083/jcb.201607043. Publisher: The Rockefeller University Press.

19. Christine J. Smoyer and Sue L. Jaspersen. Patrolling the nucleus: inner nuclear membrane-associated degradation. Current Genetics, 65(5):1099–1106, October 2019. ISSN 1432-0983. doi: 10.1007/s00294-019-00971-1.

20. Sandra Pankow, Casimir Bamberger, Diego Calzolari, Andreas Bamberger, and John R. Yates. Deep interactome profiling of membrane proteins by co-interacting protein identification technology. Nature Protocols, 11(12):2515–2528, December 2016. ISSN 1750-2799. doi: 10.1038/nprot.2016.140. Number: 12 Publisher: Nature Publishing Group.

21. Mingyu Gu, Dollie LaJoie, Opal S. Chen, Alexander von Appen, Mark S. Ladinsky, Michael J. Redd, Linda Nikolova, Pamela J. Bjorkman, Wesley I. Sundquist, Katharine S. Ullman, and Adam Frost. LEM2 recruits CHMP7 for ESCRT-mediated nuclear envelope closure in fission yeast and human cells. Proceedings of the National Academy of Sciences of the United States of America, 114(11):E2166–E2175, March 2017. ISSN 1091-6490. doi: 10.1073/pnas.1613916114.

22. Brant M. Webster, Paolo Colombi, Jens Jäger, and C. Patrick Lusk. Surveillance of Nuclear Pore Complex Assembly by ESCRT-III/Vps4. Cell, 159(2):388–401, October 2014. ISSN 0092-8674. doi: 10.1016/j.cell.2014.09.012.

23. Brant M. Webster, David J. Thaller, Jens Jäger, Sarah E. Ochmann, Sapan Borah, and C. Patrick Lusk. Chm7 and Heh1 collaborate to link nuclear pore complex quality control with nuclear envelope sealing. The EMBO Journal, page e201694574, October 2016. ISSN 0261-4189, 1460-2075. doi: 10.15252/embj.201694574.

24. I.-Ju Lee, Ema Stokasimov, Nathaniel Dempsey, Joseph M. Varberg, Etai Jacob, Sue L. Jaspersen, and David Pellman. Factors promoting nuclear envelope assembly independent of the canonical ESCRT pathway. Journal of Cell Biology, 219(6), June 2020. ISSN 0021-9525. doi: 10.1083/jcb.201908232. Publisher: The Rockefeller University Press.

25. Frank R. Neumann and Paul Nurse. Nuclear size control in fission yeast. The Journal of Cell Biology, 179(4):593–600, November 2007. ISSN 0021-9525. doi: 10.1083/jcb.200708054.

26. Jacqueline Hayles, Valerie Wood, Linda Jeffery, Kwang-Lae Hoe, Dong-Uk Kim, Han-Oh Park, Silvia Salas-Pino, Christian Heichinger, and Paul Nurse. A genome-wide resource of cell cycle and cell shape genes of fission yeast. Open Biology, 3(5):130053, May 2013. ISSN 2046-2441. doi: 10.1098/rsob.130053.

27. Helena Cantwell and Paul Nurse. A systematic genetic screen identifies essential factors involved in nuclear size control. PLOS Genetics, 15(2):e1007929, February 2019. ISSN 1553-7404. doi: 10.1371/journal.pgen.1007929.

28. Maria Makarova, Ying Gu, Jun-Song Chen, Janel Renée Beckley, Kathleen Louise Gould, and Snezhana Oliferenko. Temporal Regulation of Lipin Activity Diverged to Account for Differences in Mitotic Programs. Current Biology, 26(2):237–243, January 2016. ISSN 0960-9822. doi: 10.1016/j.cub.2015.11.061.

29. Anete Romanauska and Alwin Köhler. The Inner Nuclear Membrane Is a Metabolically Active Territory that Generates Nuclear Lipid Droplets. Cell, 174(3):700–715.e18, July 2018. ISSN 1097-4172. doi: 10.1016/j.cell.2018.05.047.

30. Iain Hagan and Mitsuhiro Yanagida. Novel potential mitotic motor protein encoded by the fission yeast cut7 + gene. Nature, 347(6293):563–566, October 1990. ISSN 1476-4687. doi: 10.1038/347563a0. Number: 6293 Publisher: Nature Publishing Group.

31. I. Hagan and M. Yanagida. The product of the spindle formation gene sad1+ associates with the fission yeast spindle pole body and is essential for viability. Journal of Cell Biology, 129(4):1033–1047, May 1995. ISSN 0021-9525. doi: 10.1083/jcb.129.4.1033. Publisher: The Rockefeller University Press.

32. Andrew J. Bestul, Zulin Yu, Jay R. Unruh, and Sue L. Jaspersen. Molecular model of fission yeast centrosome assembly determined by superresolution imaging. J Cell Biol, page jcb.201701041, June 2017. ISSN 0021-9525, 1540-8140. doi: 10.1083/jcb.201701041.

33. Rebecca K. Swartz, Elisa C. Rodriguez, and Megan C. King. A role for nuclear envelope–bridging complexes in homology-directed repair. Molecular Biology of the Cell, 25 (16):2461–2471, June 2014. ISSN 1059-1524. doi: 10.1091/mbc.e13-10-0569. Publisher: American Society for Cell Biology (mboc).

34. Sarah Wälde and Megan C. King. The KASH protein Kms2 coordinates mitotic remodeling of the spindle pole body. Journal of Cell Science, 127(Pt 16):3625–3640, August 2014. ISSN 1477-9137. doi: 10.1242/jcs.154997.

35. Alfonso Fernández-Álvarez, Cécile Bez, Eileen T. O’Toole, Mary Morphew, and Julia Promisel Cooper. Mitotic Nuclear Envelope Breakdown and Spindle Nucleation Are Controlled by Interphase Contacts between Centromeres and the Nuclear Envelope. Developmental Cell, 39(5):544–559, 2016. ISSN 1878-1551. doi: 10.1016/j.devcel.2016.10.021.

36. Yuji Chikashige, Chihiro Tsutsumi, Miho Yamane, Kasumi Okamasa, Tokuko Haraguchi, and Yasushi Hiraoka. Meiotic Proteins Bqt1 and Bqt2 Tether Telomeres to Form the Bouquet Arrangement of Chromosomes. Cell, 125(1):59–69, April 2006. ISSN 0092-8674. doi: 10.1016/j.cell.2006.01.048.

37. R. R. West, E. V. Vaisberg, R. Ding, P. Nurse, and J. R. McIntosh. cut11(+): A gene required for cell cycle-dependent spindle pole body anchoring in the nuclear envelope and bipolar spindle formation in Schizosaccharomyces pombe. Molecular Biology of the Cell, 9(10):2839–2855, October 1998. ISSN 1059-1524.

38. H. J. Chial, M. P. Rout, T. H. Giddings, and M. Winey. Saccharomyces cerevisiae Ndc1p is a shared component of nuclear pore complexes and spindle pole bodies. The Journal of Cell Biology, 143(7):1789–1800, December 1998. ISSN 0021-9525.

39. Fabrizia Stavru, Bastian B. Hülsmann, Anne Spang, Enno Hartmann, Volker C. Cordes, and Dirk Görlich. NDC1: a crucial membrane-integral nucleoporin of metazoan nuclear pore complexes. The Journal of Cell Biology, 173(4):509–519, May 2006. ISSN 0021-9525. doi: 10.1083/jcb.200601001.

40. Babett Steglich, Guillaume Filion, Bas van Steensel, and Karl Ekwall. The inner nuclear membrane proteins Man1 and Ima1 link to two different types of chromatin at the nuclear periphery in *S. pombe*. Nucleus, 3(1):77–87, January 2012. ISSN 1949-1034, 1949-1042. doi: 10.4161/nucl.18825.

41. Yasushi Hiraoka, Hiromi Maekawa, Haruhiko Asakawa, Yuji Chikashige, Tomoko Kojidani, Hiroko Osakada, Atsushi Matsuda, and Tokuko Haraguchi. Inner nuclear membrane protein Ima1 is dispensable for intranuclear positioning of centromeres: S. pombe inner nuclear membrane proteins. Genes to Cells, 16(10):1000–1011, October 2011. ISSN 13569597. doi: 10.1111/j.1365-2443.2011.01544.x.

42. Charlotta Buch, Robert Lindberg, Ricardo Figueroa, Santhosh Gudise, Evgeny Onischenko, and Einar Hallberg. An integral protein of the inner nuclear membrane localizes to the mitotic spindle in mammalian cells. J Cell Sci, 122(12):2100–2107, June 2009. ISSN 0021-9533, 1477-9137. doi: 10.1242/jcs.047373.

43. Shahid Banday, Zeenat Farooq, Romana Rashid, Ehsaan Abdullah, and Mohammad Altaf. Role of Inner Nuclear Membrane Protein Complex Lem2-Nur1 in Heterochromatic Gene Silencing. The Journal of Biological Chemistry, 291(38):20021–20029, 2016. ISSN 1083-351X. doi: 10.1074/jbc.M116.743211.

44. Ramón Ramos Barrales, Marta Forn, Paula Raluca Georgescu, Zsuzsa Sarkadi, and Sigurd Braun. Control of heterochromatin localization and silencing by the nuclear membrane protein Lem2. Genes & Development, 30(2):133–148, January 2016. ISSN 1549-5477. doi: 10.1101/gad.271288.115.

45. Yoshie Tange, Yuji Chikashige, Shinya Takahata, Kei Kawakami, Masato Higashi, Chie Mori, Tomoko Kojidani, Yasuhiro Hirano, Haruhiko Asakawa, Yota Murakami, Tokuko Haraguchi, and Yasushi Hiraoka. Inner nuclear membrane protein Lem2 augments heterochromatin formation in response to nutritional conditions. Genes to Cells: Devoted to Molecular & Cellular Mechanisms, 21(8):812–832, August 2016. ISSN 1365-2443. doi: 10.1111/gtc.12385.

46. Gerard H. Pieper, Simon Sprenger, David Teis, and Snezhana Oliferenko. ESCRT-III/Vps4 Controls Heterochromatin-Nuclear Envelope Attachments. Developmental Cell, 53(1):27–41.e6, April 2020. ISSN 1534-5807. doi: 10.1016/j.devcel.2020.01.028.

47. Yanira Gonzalez, Akira Saito, and Shelley Sazer. Fission yeast Lem2 and Man1 perform fundamental functions of the animal cell nuclear lamina. Nucleus (Austin, Tex.), 3(1):60–76, February 2012. ISSN 1949-1042. doi: 10.4161/nucl.18824.

48. Kazunori Kume, Helena Cantwell, Alana Burrell, and Paul Nurse. Nuclear membrane protein Lem2 regulates nuclear size through membrane flow. Nature Communications, 10 (1):1871, April 2019. ISSN 2041-1723. doi: 10.1038/s41467-019-09623-x.

49. Yasuhiro Hirano, Yasuha Kinugasa, Hiroko Osakada, Tomoko Shindo, Yoshino Kubota, Shinsuke Shibata, Tokuko Haraguchi, and Yasushi Hiraoka. Lem2 and Lnp1 maintain the membrane boundary between the nuclear envelope and endoplasmic reticulum. Communications Biology, 3(1):1–14, June 2020. ISSN 2399-3642. doi: 10.1038/s42003-020-0999-9. Number: 1 Publisher: Nature Publishing Group.

50. Hui-Ju Yang, Masaaki Iwamoto, Yasushi Hiraoka, and Tokuko Haraguchi. Function of nuclear membrane proteins in shaping the nuclear envelope integrity during closed mitosis. The Journal of Biochemistry, 161(6):471–477, June 2017. ISSN 0021-924X. doi: 10.1093/jb/mvx020.

51. Jamie Snider, Saranya Kittanakom, Dunja Damjanovic, Jasna Curak, Victoria Wong, and Igor Stagljar. Detecting interactions with membrane proteins using a membrane two-hybrid assay in yeast. Nature Protocols, 5(7):1281–1293, July 2010. ISSN 1750-2799. doi: 10.1038/nprot.2010.83. Number: 7 Publisher: Nature Publishing Group.

52. Jingjing Chen, Christine J. Smoyer, Brian D. Slaughter, Jay R. Unruh, and Sue L. Jaspersen. The SUN protein Mps3 controls Ndc1 distribution and function on the nuclear membrane. J Cell Biol, 204(4):523–539, February 2014. ISSN 0021-9525, 1540-8140. doi: 10.1083/jcb.201307043.

53. Igor Stagljar, Chantal Korostensky, Nils Johnsson, and Stephan te Heesen. A genetic system based on split-ubiquitin for the analysis of interactions between membrane proteins in vivo. Proceedings of the National Academy of Sciences, 95(9):5187–5192, April 1998. ISSN 0027-8424, 1091-6490. doi: 10.1073/pnas.95.9.5187. Publisher: National Academy of Sciences Section: Biological Sciences.

54. John P. Miller, Russell S. Lo, Asa Ben-Hur, Cynthia Desmarais, Igor Stagljar, William Stafford Noble, and Stanley Fields. Large-scale identification of yeast integral membrane protein interactions. Proceedings of the National Academy of Sciences, 102(34):12123–12128, August 2005. ISSN 0027-8424, 1091-6490. doi: 10.1073/pnas.0505482102.

55. Christian M. Paumi, Javier Menendez, Anthony Arnoldo, Kim Engels, Kavitha Ravee Iyer, Safia Thaminy, Oleg Georgiev, Yves Barral, Susan Michaelis, and Igor Stagljar. Mapping protein-protein interactions for the yeast ABC transporter Ycf1p by integrated split-ubiquitin membrane yeast two-hybrid analysis. Molecular Cell, 26(1):15–25, April 2007. ISSN 1097-2765. doi: 10.1016/j.molcel.2007.03.011.

56. Matthew D. McGee, Igor Stagljar, and Daniel A. Starr. KDP-1 is a nuclear envelope KASH protein required for cell-cycle progression. Journal of Cell Science, 122(16):2895–2905, August 2009. ISSN 0021-9533, 1477-9137. doi: 10.1242/jcs.051607. Publisher: The Company of Biologists Ltd Section: Research Article.

57. Jay Jin, Saranya Kittanakom, Victoria Wong, Beverly AS Reyes, Elisabeth J. Van Bockstaele, Igor Stagljar, Wade Berrettini, and Robert Levenson. Interaction of the mu-opioid receptor with GPR177 (Wntless) inhibits Wnt secretion: potential implications for opioid dependence. BMC Neuroscience, 11(1):33, March 2010. ISSN 1471-2202. doi: 10.1186/1471-2202-11-33.

58. Serge M. Gisler, Saranya Kittanakom, Daniel Fuster, Victoria Wong, Mia Bertic, Tamara Radanovic, Randy A. Hall, Heini Murer, Jürg Biber, Daniel Markovich, Orson W. Moe, and Igor Stagljar. Monitoring Protein-Protein Interactions between the Mammalian Integral Membrane Transporters and PDZ-interacting Partners Using a Modified Split-ubiquitin Membrane Yeast Two-hybrid System. Molecular & Cellular Proteomics, 7(7):1362–1377, July 2008. ISSN 1535-9476, 1535-9484. doi: 10.1074/mcp.M800079-MCP200. Publisher: American Society for Biochemistry and Molecular Biology Section: Research.

59. Tommy V. Vo, Jishnu Das, Michael J. Meyer, Nicolas A. Cordero, Nurten Akturk, Xiaomu Wei, Benjamin J. Fair, Andrew G. Degatano, Robert Fragoza, Lisa G. Liu, Akihisa Matsuyama, Michelle Trickey, Sachi Horibata, Andrew Grimson, Hiroyuki Yamano, Minoru Yoshida, Frederick P. Roth, Jeffrey A. Pleiss, Yu Xia, and Haiyuan Yu. A Proteome-wide Fission Yeast Interactome Reveals Network Evolution Principles from Yeasts to Human. Cell, 164(1):310–323, January 2016. ISSN 0092-8674. doi: 10.1016/j.cell.2015.11.037.

60. Evgeny Onischenko, Leslie H. Stanton, Alexis S. Madrid, Thomas Kieselbach, and Karsten Weis. Role of the Ndc1 interaction network in yeast nuclear pore complex assembly and maintenance. The Journal of Cell Biology, 185(3):475–491, May 2009. ISSN 0021-9525, 1540-8140. doi: 10.1083/jcb.200810030.

61. Yasuhiro Hirano, Yasuha Kinugasa, Haruhiko Asakawa, Yuji Chikashige, Chikashi Obuse, Tokuko Haraguchi, and Yasushi Hiraoka. Lem2 is retained at the nuclear envelope through its interaction with Bqt4 in fission yeast. Genes to Cells: Devoted to Molecular & Cellular Mechanisms, January 2018. ISSN 1365-2443. doi: 10.1111/gtc.12557.

62. Hani Ebrahimi, Hirohisa Masuda, Devanshi Jain, and Julia Promisel Cooper. Distinct ‘safe zones’ at the nuclear envelope ensure robust replication of heterochromatic chromosome regions. eLife, 7:e32911, May 2018. ISSN 2050-084X. doi: 10.7554/eLife.32911.

63. Karim Mekhail, Jan Seebacher, Steven P. Gygi, and Danesh Moazed. Role for perinuclear chromosome tethering in maintenance of genome stability. Nature, 456(7222):667–670, December 2008. ISSN 0028-0836. doi: 10.1038/nature07460.

64. Nahid Iglesias, Joao A. Paulo, Antonis Tatarakis, Xiaoyi Wang, Amanda L. Edwards, Natarajan V. Bhanu, Benjamin A. Garcia, Wilhelm Haas, Steven P. Gygi, and Danesh Moazed. Native Chromatin Proteomics Reveals a Role for Specific Nucleoporins in Heterochromatin Organization and Maintenance. Molecular Cell, 77(1):51–66.e8, 2020. ISSN 1097-4164. doi: 10.1016/j.molcel.2019.10.018.

65. Sigurd Braun and Ramón Ramos Barrales. Beyond Tethering and the LEM domain: MS-Cellaneous functions of the inner nuclear membrane Lem2. Nucleus (Austin, Tex.), 7(6): 523–531, November 2016. ISSN 1949-1042. doi: 10.1080/19491034.2016.1252892.

66. William T. Yewdell, Paolo Colombi, Taras Makhnevych, and C. Patrick Lusk. Lumenal interactions in nuclear pore complex assembly and stability. Molecular Biology of the Cell, 22(8):1375–1388, April 2011. ISSN 1059-1524. doi: 10.1091/mbc.E10-06-0554.

67. Bernhard Moser, José Basílio, Josef Gotzmann, Andreas Brachner, and Roland Foisner. Comparative Interactome Analysis of Emerin, MAN1 and LEM2 Reveals a Unique Role for LEM2 in Nucleotide Excision Repair. Cells, 9(2), February 2020. ISSN 2073-4409. doi: 10.3390/cells9020463.

68. Adam Frost, Marc G. Elgort, Onn Brandman, Clinton Ives, Sean R. Collins, Lakshmi Miller-Vedam, Jimena Weibezahn, Marco Y. Hein, Ina Poser, Matthias Mann, Anthony A. Hyman, and Jonathan S. Weissman. Functional Repurposing Revealed by Comparing S. pombe and S. cerevisiae Genetic Interactions. Cell, 149(6):1339–1352, June 2012. ISSN 0092-8674, 1097-4172. doi: 10.1016/j.cell.2012.04.028.

69. Santhosh Gudise, Ricardo A. Figueroa, Robert Lindberg, Veronica Larsson, and Einar Hallberg. Samp1 is functionally associated with the LINC complex and A-type lamina networks. J Cell Sci, 124(12):2077–2085, June 2011. ISSN 0021-9533, 1477-9137. doi: 10.1242/jcs.078923.

70. Joana Borrego-Pinto, Thibaud Jegou, Daniel S. Osorio, Frederic Auradé, Mátyás Gor-jánácz, Birgit Koch, Iain W. Mattaj, and Edgar R. Gomes. Samp1 is a component of TAN lines and is required for nuclear movement. J Cell Sci, 125(5):1099–1105, March 2012. ISSN 0021-9533, 1477-9137. doi: 10.1242/jcs.087049.

71. Haitong Hou, Scott P. Kallgren, and Songtao Jia. Csi1 illuminates the mechanism and function of Rabl configuration. Nucleus, 4(3):176–181, May 2013. ISSN 1949-1034, 1949-1042. doi: 10.4161/nucl.24876.

72. Paola Vacchina, Karina E. J. Tripodi, Andrea M. Escalante, and Antonio D. Uttaro. Characterization of bifunctional sphingolipid Δ4-desaturases/C4-hydroxylases of trypanosomatids by liquid chromatography-electrospray tandem mass spectrometry. Molecular and Biochemical Parasitology, 184(1):29–38, July 2012. ISSN 1872-9428. doi: 10.1016/j.molbiopara.2012.04.005.

73. Petra Sperling, Philipp Ternes, Hermann Moll, Stephan Franke, Ulrich Zähringer, and Ernst Heinz. Functional characterization of sphingolipid C4-hydroxylase genes from Arabidopsis thaliana. FEBS Letters, 494(1-2):90–94, 2001. ISSN 1873-3468. doi: 10.1016/S0014-5793(01)02332-8. _eprint: https://febs.onlinelibrary.wiley.com/doi/pdf/10.1016/S0014-5793%2801%2902332-8.

74. D. Haak, K. Gable, T. Beeler, and T. Dunn. Hydroxylation of Saccharomyces cerevisiae ceramides requires Sur2p and Scs7p. The Journal of Biological Chemistry, 272(47):29704–29710, November 1997. ISSN 0021-9258. doi: 10.1074/jbc.272.47.29704.

75. Mai Nakase, Motohiro Tani, Tomotake Morita, Hiroko K. Kitamoto, Jun Kashiwazaki, Taro Nakamura, Akira Hosomi, Naotaka Tanaka, and Kaoru Takegawa. Mannosylinositol phosphorylceramide is a major sphingolipid component and is required for proper localization of plasma-membrane proteins in Schizosaccharomyces pombe. Journal of Cell Science, 123(9):1578–1587, May 2010. ISSN 0021-9533, 1477-9137. doi: 10.1242/jcs.059139. Publisher: The Company of Biologists Ltd Section: Research Article.

76. Li-Chun Cheng, Sabyasachi Baboo, Cory Lindsay, Liza Brusman, Salvador Martinez-Bartolomé, Olga Tapia, Xi Zhang, John R. Yates, and Larry Gerace. Identification of new transmembrane proteins concentrated at the nuclear envelope using organellar pro-teomics of mesenchymal cells. Nucleus, 10(1):126–143, May 2019. ISSN 1949-1034. doi: 10.1080/19491034.2019.1618175.

77. Pamela Magini, Daphne J. Smits, Laura Vandervore, Rachel Schot, Marta Columbaro, Es-mee Kasteleijn, Mees van der Ent, Flavia Palombo, Maarten H. Lequin, Marjolein Dremmen, Marie Claire Y. de Wit, Mariasavina Severino, Maria Teresa Divizia, Pasquale Stri-ano, Natalia Ordonez-Herrera, Amal Alhashem, Ahmed Al Fares, Malak Al Ghamdi, Arndt Rolfs, Peter Bauer, Jeroen Demmers, Frans W. Verheijen, Martina Wilke, Marjon van Slegtenhorst, Peter J. van der Spek, Marco Seri, Anna C. Jansen, Rolf W. Stottmann, Robert B. Hufnagel, Robert J. Hopkin, Deema Aljeaid, Wojciech Wiszniewski, Pawel Gawl-inski, Milena Laure-Kamionowska, Fowzan S. Alkuraya, Hanah Akleh, Valentina Stanley, Damir Musaev, Joseph G. Gleeson, Maha S. Zaki, Nicola Brunetti-Pierri, Gerarda Cappuccio, Bella Davidov, Lina Basel-Salmon, Lily Bazak, Noa Ruhrman Shahar, Aida Bertoli-Avella, Ghayda M. Mirzaa, William B. Dobyns, Tommaso Pippucci, Maarten Fornerod, and Grazia M. S. Mancini. Loss of SMPD4 Causes a Developmental Disorder Characterized by Microcephaly and Congenital Arthrogryposis. The American Journal of Human Genetics, 105(4):689–705, October 2019. ISSN 0002-9297. doi: 10.1016/j.ajhg.2019.08.006.

78. Sunyoung Hwang, Jessica F. Williams, Maja Kneissig, Maria Lioudyno, Isabel Rivera, Pablo Helguera, Jorge Busciglio, Zuzana Storchova, Megan C. King, and Eduardo M. Torres. Suppressing Aneuploidy-Associated Phenotypes Improves the Fitness of Trisomy 21 Cells. Cell Reports, 29(8):2473–2488.e5, 2019. ISSN 2211-1247. doi: 10.1016/j.celrep.2019.10.059.

79. Alexandra Grippa, Laura Buxó, Gabriel Mora, Charlotta Funaya, Fatima-Zahra Idrissi, Francesco Mancuso, Raul Gomez, Júlia Muntanyà, Eduard Sabidó, and Pedro Carvalho. The seipin complex Fld1/Ldb16 stabilizes ER-lipid droplet contact sites. The Journal of Cell Biology, 211(4):829–844, November 2015. ISSN 1540-8140. doi: 10.1083/jcb.201502070.

80. Alexandre Toulmay and William A. Prinz. A conserved membrane-binding domain targets proteins to organelle contact sites. Journal of Cell Science, 125(Pt 1):49–58, January 2012. ISSN 1477-9137. doi: 10.1242/jcs.085118.

81. Li-Ka Liu, Vineet Choudhary, Alexandre Toulmay, and William A. Prinz. An inducible ER-Golgi tether facilitates ceramide transport to alleviate lipotoxicity. The Journal of Cell Biology, 216(1):131–147, January 2017. ISSN 1540-8140. doi: 10.1083/jcb.201606059.

82. Jinsil Kim, Hye-Jeong Ha, Sujin Kim, Ah-Reum Choi, Sook-Jeong Lee, Kwang-Lae Hoe, and Dong-Uk Kim. Identification of Rbd2 as a candidate protease for sterol regulatory element binding protein (SREBP) cleavage in fission yeast. Biochemical and Biophysical Research Communications, 468(4):606–610, December 2015. ISSN 1090-2104. doi: 10.1016/j.bbrc.2015.10.165.

83. Jiwon Hwang, Diedre Ribbens, Sumana Raychaudhuri, Leah Cairns, He Gu, Adam Frost, Siniša Urban, and Peter J. Espenshade. A Golgi rhomboid protease Rbd2 recruits Cdc48 to cleave yeast SREBP. The EMBO journal, 35(21):2332–2349, 2016. ISSN 1460-2075. doi: 10.15252/embj.201693923.

84. Margaret I. Kanipes, John E. Hill, and Susan A. Henry. The Schizosaccharomyces pombe cho1+ Gene Encodes a Phospholipid Methyltransferase. Genetics, 150(2):553–562, October 1998. ISSN 0016-6731, 1943-2631. Publisher: Genetics Section: Investigations.

85. Alexis S. Madrid, Joel Mancuso, W. Zacheus Cande, and Karsten Weis. The role of the integral membrane nucleoporins Ndc1p and Pom152p in nuclear pore complex assembly and function. The Journal of Cell Biology, 173(3):361–371, May 2006. ISSN 0021-9525, 1540-8140. doi: 10.1083/jcb.200506199.

86. Jörg Mansfeld, Stephan Güttinger, Lisa A. Hawryluk-Gara, Nelly Panté, Moritz Mall, Vincent Galy, Uta Haselmann, Petra Mühlhäusser, Richard W. Wozniak, Iain W. Mattaj, Ulrike Kutay, and Wolfram Antonin. The Conserved Transmembrane Nucleoporin NDC1 Is Required for Nuclear Pore Complex Assembly in Vertebrate Cells. Molecular Cell, 22(1): 93–103, April 2006. ISSN 1097-2765. doi: 10.1016/j.molcel.2006.02.015.

87. Dan Zhang and Snezhana Oliferenko. Tts1, the fission yeast homologue of the TMEM33 family, functions in NE remodeling during mitosis. Molecular Biology of the Cell, 25(19): 2970–2983, October 2014. ISSN 1059-1524, 1939-4586. doi: 10.1091/mbc.E13-12-0729.

88. T. Renee Dawson, Michelle D. Lazarus, Martin W. Hetzer, and Susan R. Wente. ER mem-brane–bending proteins are necessary for de novo nuclear pore formation. The Journal of Cell Biology, 184(5):659–675, March 2009. ISSN 0021-9525. doi: 10.1083/jcb.200806174.

89. Amanda K. Casey, T. Renee Dawson, Jingjing Chen, Jennifer M. Friederichs, Sue L. Jaspersen, and Susan R. Wente. Integrity and function of the Saccharomyces cerevisiae spindle pole body depends on connections between the membrane proteins Ndc1, Rtn1, and Yop1. Genetics, 192(2):441–455, October 2012. ISSN 1943-2631. doi: 10.1534/genetics.112.141465.

90. Amanda K. Casey, Shuliang Chen, Peter Novick, Susan Ferro-Novick, and Susan R. Wente. Nuclear pore complex integrity requires Lnp1, a regulator of cortical endoplasmic reticulum. Molecular Biology of the Cell, 26(15):2833–2844, August 2015. ISSN 1939-4586. doi: 10.1091/mbc.E15-01-0053.

91. Lisa A. Hawryluk-Gara, Melpomeni Platani, Rachel Santarella, Richard W. Wozniak, and Iain W. Mattaj. Nup53 Is Required for Nuclear Envelope and Nuclear Pore Complex Assembly. Molecular Biology of the Cell, 19(4):1753–1762, April 2008. ISSN 1059-1524. doi: 10.1091/mbc.E07-08-0820.

92. Junjie Hu, Yoko Shibata, Christiane Voss, Tom Shemesh, Zongli Li, Margaret Coughlin, Michael M. Kozlov, Tom A. Rapoport, and William A. Prinz. Membrane Proteins of the Endoplasmic Reticulum Induce High-Curvature Tubules. Science, 319(5867):1247–1250, February 2008. ISSN 0036-8075, 1095-9203. doi: 10.1126/science.1153634.

93. Junjie Hu, Yoko Shibata, Peng-Peng Zhu, Christiane Voss, Neggy Rismanchi, William A. Prinz, Tom A. Rapoport, and Craig Blackstone. A class of dynamin-like GTPases involved in the generation of the tubular ER network. Cell, 138(3):549–561, August 2009. ISSN 0092-8674. doi: 10.1016/j.cell.2009.05.025.

94. Gia K. Voeltz, William A. Prinz, Yoko Shibata, Julia M. Rist, and Tom A. Rapoport. A class of membrane proteins shaping the tubular endoplasmic reticulum. Cell, 124(3):573–586, February 2006. ISSN 0092-8674. doi: 10.1016/j.cell.2005.11.047.

95. Christiane Voss, Sujoy Lahiri, Barry P. Young, Christopher J. Loewen, and William A. Prinz. ER-shaping proteins facilitate lipid exchange between the ER and mitochondria in S. cerevisiae. Journal of Cell Science, 125(Pt 20):4791–4799, October 2012. ISSN 1477-9137. doi: 10.1242/jcs.105635.

96. Christine M. Doucet and Martin W. Hetzer. Nuclear pore biogenesis into an intact nuclear envelope. Chromosoma, 119(5):469–477, October 2010. ISSN 1432-0886. doi: 10.1007/s00412-010-0289-2.

97. Anne Chadrin, Barbara Hess, Mabel San Roman, Xavier Gatti, Bérangère Lombard, Damarys Loew, Yves Barral, Benoit Palancade, and Valérie Doye. Pom33, a novel trans-membrane nucleoporin required for proper nuclear pore complex distribution. The Journal of Cell Biology, 189(5):795–811, May 2010. ISSN 1540-8140. doi: 10.1083/jcb.200910043.

98. Takeshi Urade, Yasunori Yamamoto, Xia Zhang, Yonson Ku, and Toshiaki Sakisaka. Identification and characterization of TMEM33 as a reticulon-binding protein. The Kobe Journal of Medical Sciences, 60(3):E57–65, November 2014. ISSN 1883-0498.

99. Sue L. Jaspersen and Suman Ghosh. Nuclear envelope insertion of spindle pole bodies and nuclear pore complexes. Nucleus, 3(3):226–236, May 2012. ISSN 1949-1034. doi: 10.4161/nucl.20148.

100. Jingjing Chen, Jennifer M. Gardner, Zulin Yu, Sarah E. Smith, Sean McKinney, Brian D. Slaughter, Jay R. Unruh, and Sue L. Jaspersen. Yeast centrosome components form a noncanonical LINC complex at the nuclear envelope insertion site. J Cell Biol, page jcb.201809045, March 2019. ISSN 0021-9525, 1540-8140. doi: 10.1083/jcb.201809045.

101. Thomas Kupke, Jörg Malsam, and Elmar Schiebel. A Ternary Membrane Protein Complex Anchors the Spindle Pole Body in the Nuclear Envelope in Budding Yeast. Journal of Biological Chemistry, page jbc.M117.780601, March 2017. ISSN 0021-9258, 1083-351X. doi: 10.1074/jbc.M117.780601. Publisher: American Society for Biochemistry and Molecular Biology.

102. Judite Costa, Chuanhai Fu, V. Mohini Khare, and Phong T. Tran. csi2p modulates microtubule dynamics and organizes the bipolar spindle for chromosome segregation. Molecular Biology of the Cell, 25(24):3900–3908, December 2014. ISSN 1939-4586. doi: 10.1091/mbc.E14-09-1370.

103. I. Roussou and G. Draetta. The Schizosaccharomyces pombe casein kinase II alpha and beta subunits: evolutionary conservation and positive role of the beta subunit. Molecular and Cellular Biology, 14(1):576–586, January 1994. ISSN 0270-7306. doi: 10.1128/mcb.14.1.576.

104. Atsushi Shimada, Kohei Dohke, Mahito Sadaie, Kaori Shinmyozu, Jun-Ichi Nakayama, Takeshi Urano, and Yota Murakami. Phosphorylation of Swi6/HP1 regulates transcriptional gene silencing at heterochromatin. Genes & Development, 23(1):18–23, January 2009. ISSN 0890-9369, 1549-5477. doi: 10.1101/gad.1708009. Company: Cold Spring Harbor Laboratory Press Distributor: Cold Spring Harbor Laboratory Press Institution: Cold Spring Harbor Laboratory Press Label: Cold Spring Harbor Laboratory Press Publisher: Cold Spring Harbor Lab.

105. Julie Blyth, Vasso Makrantoni, Rachael E. Barton, Christos Spanos, Juri Rappsilber, and Adele L. Marston. Genes Important for Schizosaccharomyces pombe Meiosis Identified Through a Functional Genomics Screen. Genetics, 208(2):589–603, 2018. ISSN 1943-2631. doi: 10.1534/genetics.117.300527.

106. Midori Ohta, Masamitsu Sato, and Masayuki Yamamoto. Spindle pole body components are reorganized during fission yeast meiosis. Molecular Biology of the Cell, 23(10):1799–1811, May 2012. ISSN 1939-4586. doi: 10.1091/mbc.E11-11-0951.

107. S. Ikemoto, T. Nakamura, M. Kubo, and C. Shimoda. S. pombe sporulation-specific coiled-coil protein Spo15p is localized to the spindle pole body and essential for its modification. Journal of Cell Science, 113 (Pt 3):545–554, February 2000. ISSN 0021-9533.

108. F. Miki, A. Kurabayashi, Y. Tange, K. Okazaki, M. Shimanuki, and O. Niwa. Two-hybrid search for proteins that interact with Sad1 and Kms1, two membrane-bound components of the spindle pole body in fission yeast. Molecular genetics and genomics: MGG, 270(6): 449–461, January 2004. ISSN 1617-4615. doi: 10.1007/s00438-003-0938-8.

109. Ayumu Yamamoto, Robert R. West, J. Richard McIntosh, and Yasushi Hiraoka. A Cytoplasmic Dynein Heavy Chain Is Required for Oscillatory Nuclear Movement of Meiotic Prophase and Efficient Meiotic Recombination in Fission Yeast. The Journal of Cell Biology, 145(6):1233–1250, June 1999. ISSN 0021-9525.

110. Kazunori Tomita and Julia Promisel Cooper. The Telomere Bouquet Controls the Meiotic Spindle. Cell, 130(1):113–126, July 2007. ISSN 0092-8674, 1097-4172. doi: 10.1016/j.cell.2007.05.024. Publisher: Elsevier.

111. Teresa Niccoli, Akira Yamashita, Paul Nurse, and Masayuki Yamamoto. The p150-Glued Ssm4p regulates microtubular dynamics and nuclear movement in fission yeast. Journal of Cell Science, 117(Pt 23):5543–5556, November 2004. ISSN 0021-9533. doi: 10.1242/jcs.01475.

112. Futaba Miki, Koei Okazaki, Mizuki Shimanuki, Ayumu Yamamoto, Yasushi Hiraoka, and Osami Niwa. The 14-kDa dynein light chain-family protein Dlc1 is required for regular oscillatory nuclear movement and efficient recombination during meiotic prophase in fission yeast. Molecular Biology of the Cell, 13(3):930–946, March 2002. ISSN 1059-1524. doi: 10.1091/mbc.01-11-0543.

113. Thibault Courtheoux, Guillaume Gay, Céline Reyes, Sherilyn Goldstone, Yannick Gachet, and Sylvie Tournier. Dynein participates in chromosome segregation in fission yeast. Biology of the Cell, 99(11):627–637, November 2007. ISSN 1768-322X. doi: 10.1042/BC20070047.

114. Luther Davis and Gerald R. Smith. Dynein promotes achiasmate segregation in Schizosaccharomyces pombe. Genetics, 170(2):581–590, June 2005. ISSN 0016-6731. doi: 10.1534/genetics.104.040253.

115. Mariola R. Chacón, Petrina Delivani, and Iva M. Tolić. Meiotic Nuclear Oscillations Are Necessary to Avoid Excessive Chromosome Associations. Cell Reports, 17(6):1632–1645, 2016. ISSN 2211-1247. doi: 10.1016/j.celrep.2016.10.014.

116. André Koch, Karsten Krug, Stuart Pengelley, Boris Macek, and Silke Hauf. Mitotic substrates of the kinase aurora with roles in chromatin regulation identified through quantitative phosphoproteomics of fission yeast. Science Signaling, 4(179):rs6, June 2011. ISSN 1937-9145. doi: 10.1126/scisignal.2001588.

117. Matthew P. Swaffer, Andrew W. Jones, Helen R. Flynn, Ambrosius P. Snijders, and Paul Nurse. Quantitative Phosphoproteomics Reveals the Signaling Dynamics of Cell-Cycle Kinases in the Fission Yeast Schizosaccharomyces pombe. Cell Reports, 24(2):503–514, 2018. ISSN 2211-1247. doi: 10.1016/j.celrep.2018.06.036.

118. R. Daniel Gietz and Robin A. Woods. Transformation of yeast by lithium acetate/single-stranded carrier DNA/polyethylene glycol method. In Christine Guthrie and Gerald R. Fink, editors, Methods in Enzymology, volume 350 of Guide to Yeast Genetics and Molecular and Cell Biology - Part B, pages 87–96. Academic Press, January 2002. doi: 10.1016/S0076-6879(02)50957-5.

119. Johanne M. Murray, Adam T. Watson, and Antony M. Carr. Transformation of Schizosaccharomyces pombe: Lithium Acetate/ Dimethyl Sulfoxide Procedure. Cold Spring Harbor Protocols, 2016(4):pdb.prot090969, April 2016. ISSN 1940-3402, 1559-6095. doi: 10.1101/pdb.prot090969. Publisher: Cold Spring Harbor Laboratory Press.

120. Igor Stagljar and Stanley Fields. Analysis of membrane protein interactions using yeast-based technologies. Trends in Biochemical Sciences, 27(11):559–563, November 2002. ISSN 0968-0004.

121. Safia Thaminy, John Miller, and Igor Stagljar. The split-ubiquitin membrane-based yeast two-hybrid system. Methods in Molecular Biology (Clifton, N.J.), 261:297–312, 2004. ISSN 1064-3745. doi: 10.1385/1-59259-762-9:297.

122. RStudio Team. RStudio: Integrated Development Environment for R, 2020.

123. Omar Wagih, Matej Usaj, Anastasia Baryshnikova, Benjamin VanderSluis, Elena Kuzmin, Michael Costanzo, Chad L. Myers, Brenda J. Andrews, Charles M. Boone, and Leopold Parts. SGAtools: one-stop analysis and visualization of array-based genetic interaction screens. Nucleic Acids Research, 41(W1):W591–W596, July 2013. ISSN 0305-1048. doi: 10.1093/nar/gkt400. Publisher: Oxford Academic.

124. Omar Wagih and Leopold Parts. gitter: a robust and accurate method for quantification of colony sizes from plate images. G3 (Bethesda, Md.), 4(3):547–552, March 2014. ISSN 2160-1836. doi: 10.1534/g3.113.009431.

125. Elizabeth I. Boyle, Shuai Weng, Jeremy Gollub, Heng Jin, David Botstein, J. Michael Cherry, and Gavin Sherlock. GO::TermFinder–open source software for accessing Gene Ontology information and finding significantly enriched Gene Ontology terms associated with a list of genes. Bioinformatics (Oxford, England), 20(18):3710–3715, December 2004. ISSN 1367-4803. doi: 10.1093/bioinformatics/bth456.

126. Caroline A. Schneider, Wayne S. Rasband, and Kevin W. Eliceiri. NIH Image to ImageJ: 25 years of image analysis. Nature Methods, 9(7):671–675, July 2012. ISSN 1548-7105.

